# Non-coding variants impact *cis*-regulatory coordination in a cell type-specific manner

**DOI:** 10.1101/2023.10.11.561870

**Authors:** Olga Pushkarev, Guido van Mierlo, Judith F. Kribelbauer, Wouter Saelens, Vincent Gardeux, Bart Deplancke

## Abstract

**BACKGROUND:** Interactions among *cis*-regulatory elements (CREs) play a crucial role in gene regulation. Various approaches have been developed to map these interactions genome-wide, including those relying on interindividual epigenomic variation to identify groups of covariable regulatory elements, referred to as chromatin modules (CMs). While CM mapping allows to investigate the relationship between chromatin modularity and gene expression, the computational principles used for CM identification vary in their application and outcomes.

**RESULTS:** We comprehensively evaluate and streamline existing CM mapping tools and present guidelines for optimal utilization of epigenome data from a diverse population of individuals to assess regulatory coordination across the human genome. We showcase the effectiveness of our recommended practices by analysing distinct cell types and demonstrate cell type-specificity of CRE interactions in CMs and their relevance for gene expression. Integration of genotype information revealed that many non-coding disease-associated variants affect the activity of CMs in a cell type-specific manner by affecting the binding of cell type-specific transcription factors. We provide example cases that illustrate in detail how CMs can be used to deconstruct GWAS loci, understand variable expression of cell surface receptors in immune cells and reveal how genetic variation can impact the expression of prognostic markers in chronic lymphocytic leukaemia.

**CONCLUSIONS:** Our study presents an optimal strategy for CM mapping, and reveals how CMs capture the coordination of CREs and its impact on gene expression. Non-coding genetic variants can disrupt this coordination, and we highlight how this may lead to disease predisposition in a cell type-specific manner.

## Introduction

The genome inside the nucleus is spatially organized with every chromosome occupying its own territory. Each chromosome is coarsely divided into active (A) and inactive (B) compartments [1], and Topologically Associating Domains (TADs) [2,3]. Within TADs, fine-grained chromatin organization is established through interactions of non-coding, *cis*-regulatory elements (CREs; here defined as enhancers and promoters). Such CRE interactions are critical for gene regulation, as most promoters alone possess low intrinsic power for regulating gene expression [4], and active genes interact on average with 2-4 enhancers [5,6]. Depending on the context, perturbation of one or multiple enhancers can be sufficient to largely attenuate gene expression through disruption of regulatory CRE interactions [7–9].

CREs can be accurately mapped using epigenome profiling assays such as ATAC-seq and ChIP-seq, and stratified into different categories according to their regulatory properties (enhancer vs promoter, weak vs strong etc.) on a per cell type basis [10]. Further epigenome-based studies on cohorts of non-related individuals in single cell types revealed that not every CRE harbours equal chromatin accessibility or activity across individuals [11]. Downstream association studies pairing this activity to commonly occurring genetic variants allowed to identify loci (i.e., Quantitative Trait Loci; QTLs) that impact either CRE accessibility due to disruption of transcription factor (TF) binding site(s) or histone modification deposition [12,13]. Such a ‘population epigenomics’ approach has been used to link genetic variants to the activity of CREs and gene expression in a range of cell types and conditions [14–16]. While highly valuable, these studies mostly focused on individual CREs and thus did not explicitly assess how CREs interact to regulate the expression of nearby genes.

Chromosome conformation capture (3C) as well as microscopy-based approaches allow linking enhancers and promoters into units of functionally collaborating CREs, although this can be challenging in terms of resolution and throughput[17]. An orthogonal approach is to perform epigenome analyses on many different genotypes, enabling the identification of co-variable CREs. This in turn allows the grouping of CREs based on their interindividual variability into coordinated, epigenomic hubs within TADs, which are referred to as chromatin modules (CMs [17]). Such an approach allows the mapping of CRE coordination in a genome-wide manner and, as compared to Hi-C and microscopy assays, provides direct epigenomic readouts of CRE activity and therefore allows revealing the impact of genetic variation on epigenome regulation. Nevertheless, the methods for mapping of CMs are diverse in terms of underlying statistical principles (varying from hierarchical clustering of interindividual peak correlations to Bayesian modelling [17–20]), required input data and format and interpretation of output data. Effective implementation of these approaches requires a profound understanding of underlying computational techniques, including data standardization and experience with handling different data modalities, and accurate interpretation of the results. Moreover, broad application of these methods is hindered due to the lack of streamlined pipelines and associated methodological guidelines.

To address these challenges, we evaluated the existing approaches for CM mapping in terms of methodological and CM characteristics. We assessed the (dis)advantages of each of the methods and provide guidelines related to evaluation criteria, positive controls and data and sample size. We provide executable code that will allow each researcher to easily execute each of the workflows on their own data, leading to standardized data output formats and convenient interpretation. Next, we leveraged our method benchmarking efforts to perform CM mapping in six different cell types, revealing that regulatory coordination, captured with CMs, is organized in a cell type-specific manner, with cell type-specific CREs embedded in CMs being hallmarked by binding sites for lineage-specific TFs. Through association with the underlying genotypes and focusing on distinct immune cell types, we identified multiple (autoimmune) disease-associated genetic variants that impact CM activity. These analyses complement canonical chromatin QTL mapping efforts by providing regulatory insights into how non-coding variants contribute to disease by influencing gene regulation in a cell type-specific manner.

## Results

### Comparative mapping of CMs across individuals with bulk epigenome data

Chromatin modules (CMs) represent genomic regions where CREs display co-variable activity for active histone marks such as H3K4me1 and H3K27ac [17] (**Fig 1a**). CMs can be mapped using correlation-based approaches that quantify epigenome signal co-variability across individuals to group CREs into modules using either graph-based community detection ([18], which we will refer to as VCMtools [21])) or hierarchical clustering (Clomics [19]). A complementary strategy (PHM [20]) uses interindividual variation to infer interactions between peak pairs conditioned on genetic variation (**Fig 1b**). So far, there has been no systematic comparison of existing techniques for CM mapping. To address this, we downloaded available H3K4me1 and H3K27ac ChIP-seq data for 317 lymphoblastoid cell lines (LCLs [19]) together with respective genotype data[19]. We performed peak calling on all chromosomes for H3K4me1 (218’542 peaks) and H3K27ac (127’060 peaks). We mapped CMs using VCMtools, Clomics and PHM using the smallest chromosome 22 to evaluate several aspects of computational performance (i.e., elapsed time, memory consumption), and CM-related quantitative outputs (i.e., the number of mapped CMs, median module length, and coefficient of variation of CM length) across various sample sizes (**Fig 1b, S1.1a**; Methods). Execution-wise, Clomics was the most user-friendly tool (i.e., required least input data formatting and was simple to use), the fastest (in terms of elapsed time) and the most efficient (in terms of RAM usage), followed by VCMtools and PHM (**Fig S1.1b-c**). Overall, Clomics also produced the largest number of CMs yielding two to three times more modules as compared to VCMtools and PHM (**Fig S1.1d**). The number of identified CMs and genetic associations scaled with the number of individuals and saturated at ∼175 (Clomics and VCMtools) or ∼250 (PHM) individuals, even though features such as median CM size remained relatively stable from 75 individuals onwards (**Fig S1.1d-h**). We also devised a strategy to quantify the reproducibility of mapped CMs in terms of included CREs using the different number of subsampled individuals (**Methods**). This revealed higher average reproducibility scores with larger cohort sizes, with the best average reproducibility scores achieved for >175 (Clomics / VCMtools) or 250 (PHM) samples. These analyses highlight that CM identification robustness scales with the number of included individuals, only increases marginally when more than 250 individuals are included, and requires a minimum of ∼50-75 individuals. The lower number of detected CMs with PHM compared to correlation-based approaches can be atributed to PHM identifying only those associations that are linked to an underlying, putatively causal variant. In contrast, correlation-based approaches do not rely on genotype data and are thus less constrained in terms of selection of covariable CREs.

**Figure 1.**
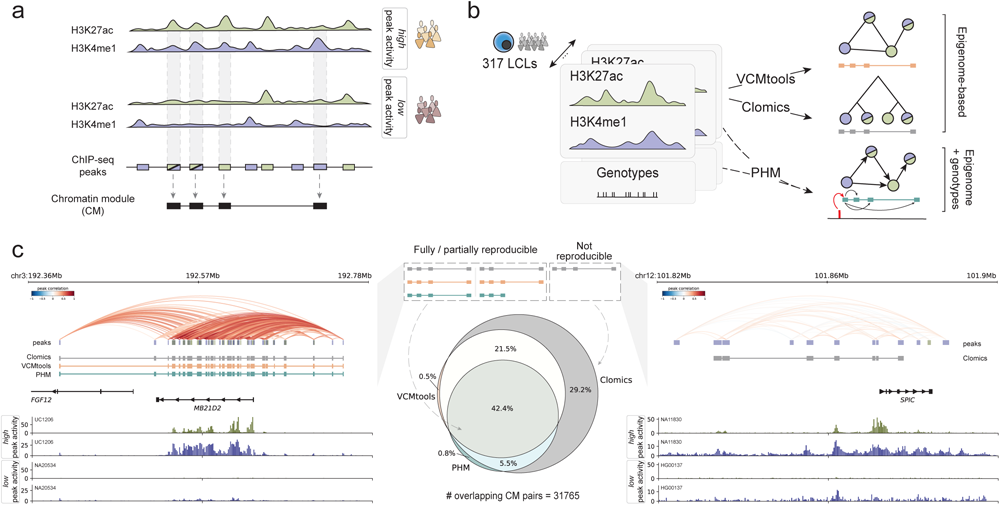
Systematic mapping of chromatin modules (CMs) in LCLs. **a.** Schematic representation of ChIP-seq profiles (H3K27ac in green, H3K4me1 in purple) for individuals with differential chromatin activity at the loci. ChIP-seq peaks are shown with purple and green rectangles, where color indicates different histone modifications. Only co-variable peaks are used to define a CM, which is depicted with black rectangles at the bottom of the panel. **b.** Schematic representation of the pipeline. We collected available H3K27ac and H3K4me1 ChIP-seq data for LCLs and associated genotypes for 317 individuals. We then used the data to map covariable regions with three approaches: correlation-based approaches (VCMtools, Clomics) that depend only on the epigenome data for CM mapping, and a Bayesian hierarchical method (PHM) that requires genotype information in addition to epigenome data. **c.** The depicted Venn diagram shows the percentage of overlapping CMs by at least one base pair across different methods. ***Left panel***: example of the most reproducible CMs across three methods in the *MD21D2* gene locus. The top track represents peak-to-peak correlations in the locus. The tracks below show CMs mapped with Clomics (gray), VCMtools (orange) and PHM (green). The bottom tracks show ChIP-seq tracks for two individuals with the most differential CM signal (H3K27ac in dark green, H3K4me1 in dark blue). ***Right panel***: example of the Clomics-specific CM (in gray) in the *SPIC* gene locus. The top track represents peak-to-peak correlations, the bottom tracks show ChIP-seq tracks for two individuals with the most differential CM signal.

Next, we identified CMs on all chromosomes with VCMtools (n=9071), Clomics (n=18633) and PHM (n=5299) (**Fig S1.1i; Table S1**). Across all overlapping CMs in pairwise method comparisons, 42.4% of all CMs were at least partially identified using all methods (Venn diagram, **Fig 1c**), with the largest average similarity between the correlation-based approaches Clomics and VCMtools (**Fig S1.2a**). A representative example of a highly reproducible CM comprises the *MB21D2* locus that was previously shown to display high levels of *cis*-regulatory coordination [20,22], displaying high or very low ChIP-seq signal in the CM-embedded regions (**Fig 1c**, left). A large proportion of CMs was identified only using Clomics (∼29%), of which a representative example is the *SPIC* locus. Although CM peaks in the *SPIC* locus showed lower average interindividual correlation (cor=0.18) compared to the CM around *MB21D2* (cor=0.58), the normalized ChIP-seq signal clearly indicates co-presence or co-absence of CM-embedded peaks (**Fig 1c**, right). This indicates that Clomics has higher power in conditions with more subtle variation in co-variable chromatin activity (**Fig S1.1j**). This is likely due to the adaptive background-aware thresholding used in Clomics that allows to account for local (1 Mb scale) differences in background correlations [19], as opposed to universal p-value thresholding implemented in VCMtools [18].

In general, the methods showed high concordance with respect to previously reported CM characteristics [17–20], such as CM lengths (median 8.6-31.7k) and sizes (median 2-4 peaks) (**Fig S1.1i**), localization (63.9-65.9% CMs in A (active) and 25.4-27.5% CMs in B (inactive) compartments, 71.3-76.8% CMs within TAD boundaries (**Fig S1.2b**), enrichment in 3D interactions between CM-embedded CREs (Hi-C 500bp **Fig S1.2c**) and active chromatin states (**Fig S1.2d-f**) (see **Methods** for details). Finally, genes overlapped by CMs (20-50% among 26362 protein coding genes and lincRNAs; 75-80% of CMs overlapped a gene) (**Fig S1.2g-h**) showed higher standard deviation and average expression compared to genes that did not overlap with a CM (**Fig S1.3a-c**), and enrichment in B cell and immunity-related terms (**Fig S1.3d**).

Based on the performed analysis, we suggest to opt for Clomics over VCMtools among correlation-based approaches. Even though there is a high overlap between the methods, Clomics stands out by its ease of use and higher sensitivity towards identifying subtle yet putatively relevant chromatin co-variation. PHM provides directional insights into CRE communication, yet requires large sample cohorts and significant computational resources. Therefore, we suggest using PHM as an auxiliary approach for interpretation of CMs mapped with Clomics, which we will hereafter use for all downstream analyses.

### Cell type specificity of interindividual variation

Understanding regulatory variation among individuals in different cell types and states is crucial both from a fundamental and translational perspective [23–25]. To explore the extent of cell type specificity of chromatin activity variation among individuals, we extended the LCL dataset with downloaded ChIP-seq (H3K27ac, H3K4me1) and genotype data for hundreds of individuals resulting in epigenomic variation coverage in five cell types: LCLs (n=317), Fibroblasts (FIB, n=78), Monocytes (n=172), Neutrophils (n=164) and T cells (n=93) [19,26]. To facilitate the interpretation of mapped co-variable CREs and ensure consistency in downstream analyses, we remapped ChIP-seq peaks to obtain a universal set of peaks for H3K27ac and H3K4me1 in all cell types (**Fig 2a, S2.1a,** Methods), and mapped CMs using Clomics in FIB (n=10158), Monocytes (n=9002), Neutrophils (n=6604) and T cells (n=4841) (**Fig S2.1b,c; Table S2**). We first assessed cell type similarity based on mapped CMs across cell types. The highest average similarity was observed for CMs mapped in Monocytes and Neutrophils, while Fibroblasts tended to be the most distant cell type from the other ones, as expected based on its cell lineage (**Fig S2.1d**). We used pairwise cell type comparisons to coarsely categorize CMs into: 1) universal CMs (similarity score > 0.7), 2) partially overlapping CMs (0 < similarity score ≤ 0.7), and 3) cell type-specific CMs (similarity score = 0). This showed that the majority of CMs are cell type-specific while some cell types, such as Monocytes and Neutrophils, have a relatively higher proportion of similar CMs (**Fig S2.1e-i**). Representative loci for each category are shown in **Fig 2b**. This high degree of cell type specificity is also reflected in the association between CM activity, captured as the first principal component of the PCA performed on the CM peak count matrix (aCM [18]), and gene expression on these representative loci (**Fig S2.2a-c**). Together, our results demonstrate the prevalent lineage-and cell type specificity of the regulatory landscapes that we captured in the form of CMs.

**Figure 2.**
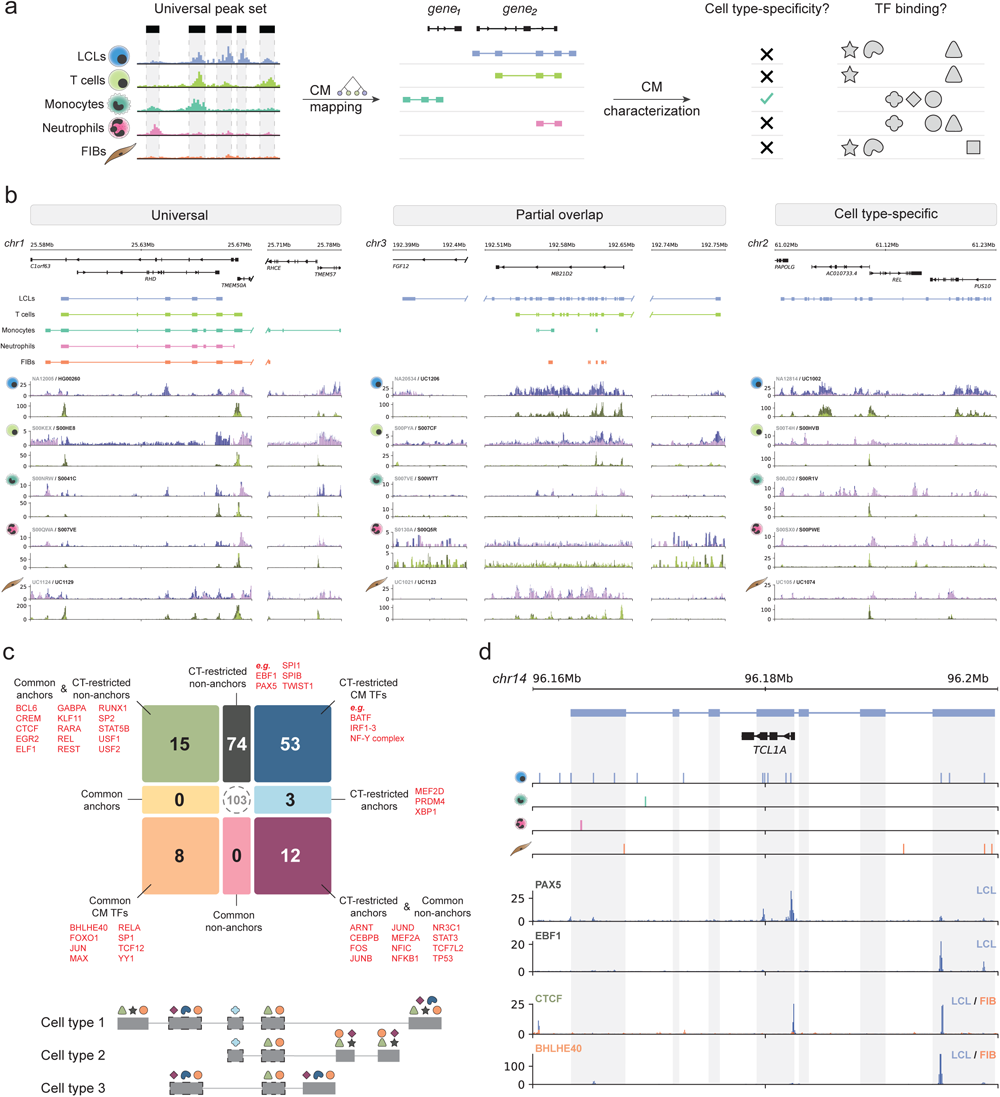
Cell type specificity of regulatory variation and TF binding in CMs. **a.** Schematic representation of the pipeline. ChIP-seq and genotype data were collected for hundreds of individuals for five cell types, namely LCLs, Monocytes, Neutrophils, T cells and Fibroblasts (FIB) [19,26]. The collected ChIP-seq data was processed in a standardized way to obtain a count matrix for a universal peak set (see **Methods** for details), which was used for CM mapping. The downstream analysis included evaluation of cell type specificity of regulatory variation captured in the form of CMs, and quantification of TFBS enrichment in specific CM peak types. **b. *From left to right***: 1. Example of a “universal” CM, here in the *RHD* gene locus where covariable peaks are present in all cell types. 2. Example of a lineage-specific CM in LCLs (blue) and T cells (light green), spanning the *MB21D2* gene. 3. Example of an LCL-specific CM in the *REL* locus. The tracks below correspond to ChIP-seq profiles of H3K27ac and H3K4me1 for two individuals, one with the highest (dark blue (H3K4me1) and dark green (H3K27ac)) and one with the lowest (magenta (H3K4me1) and lime color (H3K27ac)) CM activity. The individuals at the extremes of aCM score were selected on the basis of the largest CM in the locus by calculating pseudo-aCM score in cell types with smaller/absent CMs. **c**. TF classification according to TFBS enrichments in distinct categories stratified into anchor versus non-anchor CM CREs and cell type-restricted versus cell type-common CM CREs based on all pair-wise comparisons together (see **Methods** for details). **Bottom**: schematic representation of TFBS for different categories of TFs across partially overlapping CMs. **d**. Example of an LCL-specific CM (in blue) in the *TCL1A* locus. Tracks below indicate TFBSs (vertical lines) for cell type-enriched TFs per cell type (***from top to bottom***: LCL: ATF2 (n=1), EBF1 (n=11), KLF1 (n=1), PAX5 (n=5), Monocytes: KLF4 (n=1), Neutrophils: KLF5 (n=2), FIB: ERG (n=2), NR2F2 (n=1), TFEB (n=1)). No T cell-specific TFBS were found in the locus. The bottom tracks show ENCODE ChIP-seq profiles for PAX5 and EBF1 binding in LCLs, CTCF and BHLHE40 tracks for LCLs (blue) and FIB (orange).

### CM formation appears driven by functionally distinct groups of TFs

To identify candidate TFs associated with CREs embedded within CMs, we first performed differential peak-based TF binding site (TFBSs, which represent the genomic locations of TF motifs matching TF binding sites as determined using ChIP-seq [27]) enrichment analysis between CREs embedded within CMs and non-CM CREs for each cell type (**Fig S2.2d**). As non-CM reference regions, we created a set of ‘simulated’ CMs from the non-CM CREs that were size-, distance-, ChIP signal strength-, and GC-matched to the mapped CMs (**Methods**). The TFBS analyses revealed that CREs in CMs are significantly enriched for TFBSs of known cell type-specific TFs, e.g., EBF1 in LCLs [28] and CEBPA/B in Monocytes and Neutrophils [29] (**Fig S2.2e-i**; **Methods**), with larger similarity between myeloid (Monocytes and Neutrophils) and lymphoid (LCLs and T-cells) cell types (**Fig S2.2j**). Since CMs were mapped based on a universal peak set across cell types, this indicates that CM CREs are enriched for cell type-specific TFBSs.

To further characterize the cell type specificity of the CM-embedded CREs, we aimed to compare the TFBS enrichment between cell types as well as assess the regulatory activity of the CREs. To determine the later, we used available STARR-seq data from the GM12878 LCL cell line, which provides episomal CRE activity information [30]. We then performed pairwise comparisons between LCLs and every other cell type to stratify the CM-embedded CREs into those shared between the compared pair of cell types (i.e. CREs that we will refer to as ‘anchors’) and those specific to one of the cell types (i.e. CREs that we will refer to as “non-anchors”) (**Fig 2.3a**). For all pairwise comparisons of LCLs to other cell types, we observed significantly higher activity at anchor CREs, with the lowest average activity at non-anchors of non-LCL cell types (**Fig S2.3b**). This could indicate that CM CREs that are shared between cell types (anchor CREs) may have higher regulatory potential than non-anchor CREs, which themselves may have more secondary support functions as driven by cell type-specific TFs [31–33]. To test this hypothesis, we first aimed to assess which TFBSs are enriched in non-anchor CREs in a cell type-dependent manner, which revealed the enrichment of TFs with well-known functions in the respective cell types, such as KLF4 in Monocytes [34], KLF5 in Neutrophils [35], the pioneer B-cell TF EBF1 in LCLs [28] and TWIST1 in FIBs [36] (**Fig S2.3c-d**). These findings suggest that the TFBS enrichment that we observed when comparing the set of reference versus simulated CMs (**Fig S2.2e-i**; **Methods**) is driven by cell type-specific TFs that bind to non-anchor elements.

As a next step, we included the anchor CREs in our pairwise TFBS enrichment analyses to define if non-anchor CRE-enriched TFBSs can also be detected in anchor CREs and if this depends on the assayed cell types. To do so, we specifically focused on the anchor CM CREs and contrasted these with a collection of non-anchor CREs between each cell type pair. Together with the comparison of non-anchor CREs, this allowed us to broadly categorize TFs into several groups based on 1) whether enrichment of their respective BSs in CMs is specific to either anchors, non-anchors or both (with the latter representing “general” CM TFs), and 2) if this BS enrichment is specific to one or few cell types (i.e. “cell type restricted”), or if this enrichment is present in all pairwise cell type comparisons (‘common’) (**Fig 2c, S2.3e**).

We identified a total of 165 TFs that are enriched in anchor and/or non-anchor CREs (**Fig 2c; Table S4**). The TFBSs for a total of 23 TFs were always enriched at anchors. These TFs include more universally-expressed regulators such as USF1/2 and SP1/2, as well as proteins involved in regulating 3D chromatin organization such as CTCF and YY1 [3,37,38]. However, we found that the large majority of these 165 TFs (n=130) have BSs that are enriched in a cell type-restricted fashion in non-anchor or all CM CREs.

Altogether, these analyses suggest that CMs consist of both common regions (anchors) with higher regulatory activity that are preferentially bound by universally expressed TFs as well as CREs that are more cell type-specific and that are bound by TFs relevant to the respective cell type(s). A notable example that illustrates how CM formation may be driven by distinct classes of TFs is the B-cell-relevant *TCL1A* locus [39] where the gene is specifically expressed in B cells (**Fig S2.4e**). The *TCL1A* gene locus is enriched for binding sites of LCL-enriched TFs in the CM body, especially at the CM peaks, as compared to other cell types. This is exemplified by binding of cell type-specific TFs such as EBF1 and PAX5, as well as LCL-specific binding of CTCF and BHLHE40, at the locus (**Fig 2e, Fig S2.4f**). Together, our analyses indicate putatively divergent functional roles of TFs in the context of CM formation.

### Chromatin modules capture CREs associated with gene expression

Having identified TFs that are enriched within CMs and that as such may contribute to their establishment, we next aimed to conceptually assess how CMs can aid in understanding gene regulation (as gene expression (RNA-seq) data is not used for the mapping of CMs). It is conceivable that CMs offer several advantages compared to single CREs, since i) they group CREs into collaborating hubs, ii) the activity state of the CM (aCM) can be derived which reflects the compound activity of all CM-embedded CREs and thus that of the locus, and iii) CREs are linked to genes based on co-activity profiles and thus do not require 3D information for this. We first aimed to assess to what extent the aCM score is associated to gene expression relative to that of individual CREs (**Fig 3a**). Specifically, for each gene, we correlated its expression with 1) the height of the peak closest to the TSS but not part of the respective CM, 2) the height of the CM-embedded peak closest to the TSS, and 3) the activity score of the CM closest to the TSS. This revealed that in all conditions, the CREs closest to the TSS have the highest correlation to gene expression. However, it also revealed that CM CREs show higher correlations with expression of the closest gene at larger genomic distances compared to non-CM-embedded CREs and the group of any peaks (without distinguishing between CM and non-CM-embedded peaks) (**Fig 3b, S3.1a-d**). This can be also observed in the gain of significant associations to gene expression when using aCM scores (**Fig S3.1e-i**). In addition, absolute correlation values between aCM scores and expression of the closest genes were on average higher than absolute correlation values between the heights of individual CM peaks and expression of the closest genes, largely independent of which CM peak was assessed (**Fig S3.1j-n**). Together, by using multiple co-varying CREs aggregated into this aCM score, the set of candidate genes for which their respective expression correlates with chromatin state activity can be expanded by at least two-fold (**FigS3.2a-e**).

**Figure 3.**
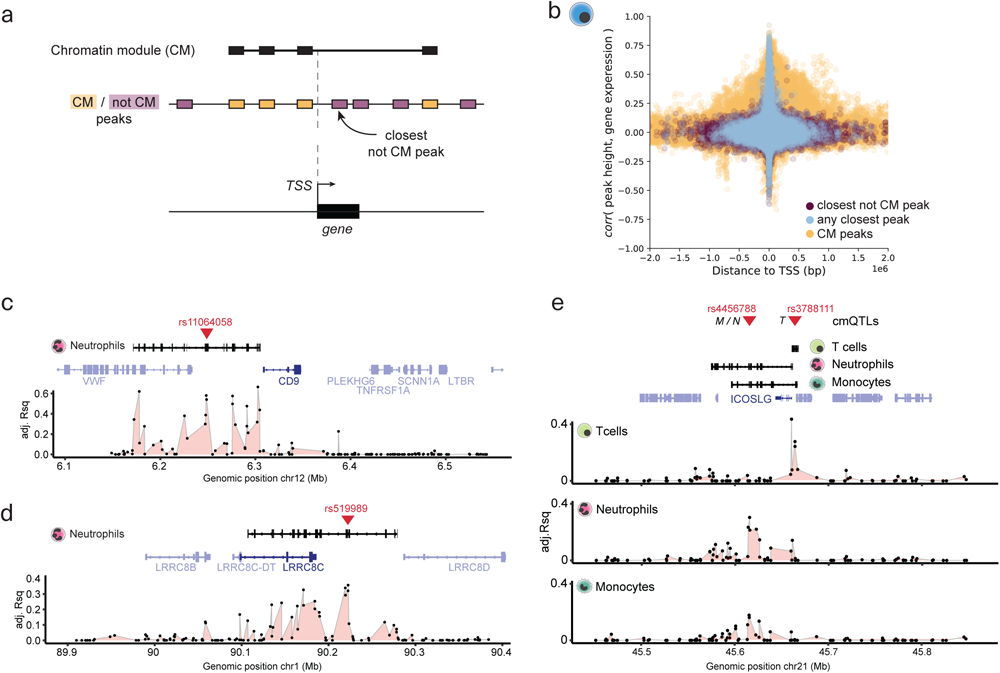
Chromatin modules capture CREs associated with gene expression. **a.** Schematic representation of how we identified CM and non-CM peaks closest to the gene transcription start site (TSS) to test the strength of association of either peak height with closest gene expression or aCM score with closest gene expression. Correlations were computed between peak heights and closest gene expression and between aCM scores and closest gene expression. **b.** Distance to the closest gene TSS from the peak center versus correlation between the peak height and the closest gene expression. The colors indicate different peak categories, where “any peak” group corresponds to the case when the closest peak is selected irrespective of its annotation as CM peak or non-CM peak. **c-e.** Examples of CMs spanning genes in various cell types having a cmQTL (red triangles). The tracks below CMs and genes show the association strength (adjusted R^2^ (Rsq) of the linear regression) between every peak in the locus and expression of the gene highlighted in dark blue.

CMs are detected by leveraging interindividual variation, so it is conceivable that genetic variation contributes to the activity of at least a portion of CMs [18,19], and through the CM affects gene expression. We therefore mapped not only histone mark (H3K4me1 and H3K27ac) QTLs (hQTLs), but also variants that affect the activity of CMs (cmQTLs; **Table S3**). While the total number of hQTLs is higher, QTLs could be mapped for a higher proportion of CMs compared to histone marks (∼10% compared to ∼40-50% for CMs; **Fig S4.1a-b**). We observed that ∼60-70% of cmQTLs are also hQTLs or in LD (R2 > 0.8) with hQTLs, indicating that in a large number of cases, variants disrupt a CM by disrupting histone mark enrichment at one or multiple CREs (**Fig S4.1c**), and they do so in a largely cell type-specific manner (**Fig S4.1d**). Binning of the interindividual peak signal variability further showed that CREs with higher interindividual variation in peak signal are more likely to be associated to a QTL and/or embedded in a CM, and that CM mapping allows to capture almost twice as many variable peaks compared to conventional hQTLs (**Fig S4.1e**). This suggests that many CMs capture regions with variable peaks that are not impacted by a focal genetic variant, but rather by another peak within the same CM. This is consistent with observations based on *in silico* approaches which suggest that frequently one or two CREs may act as the ‘dominant’ or ‘lead’ element(s) resulting in increased chromatin activity at the lead element itself as well as at nearby CREs [20]. Given that approximately half of the CMs can be linked to a candidate causal variant (**Fig S4.1a**), it is conceivable that such QTLs disrupt the lead CRE in a locus leading to both local and distal changes in the chromatin landscape. As is evident from the examples highlighted in **Fig 3c,d** and **Fig S3.2f-j**, there are generally two or three CREs at which the ChIP-signal is strongest associated with expression of the nearby gene, in line with previous observations using CRISPR interference [40,41]. The associated cmQTLs localize in one of the peaks with the strongest association (Adj. R^2^ of linear regression between peak height and gene expression; **Fig 3c,d; S3.2f-i**, red triangles), which therefore represents a putative ‘lead’ regulatory element in a genomic region. For example, in the *CD9* locus in Neutrophils, there are six principal regions that correlate with *CD9* expression (Adj. R^2^ > 0.4 & FDR < 0.05 in linear regression) (**Fig 3c**). All these elements are embedded into a CM, highlighting that these CREs covary between individuals in terms of epigenomic signal, which then influences *CD9* expression. The genetic variant with the strongest association to CM activity localizes in one of these peaks ∼50 kb upstream of the *CD9* promoter, indicating that this is the putative lead CRE for the CM. Another example comprises the *LRRC8C* locus, where we identified a cmQTL (rs519989) for the CM in Neutrophils localized in an intergenic enhancer ∼25 kb upstream of *LRRC8C* gene (**Fig 3d**). The rs519989 variant was recently shown to impact PU.1 binding to this enhancer, leading to reduced enhancer-promoter connectivity and lower expression of *LRRC8C* [42]. Together, this illustrates how CM mapping can aid in identifying causal variants and linking them to genes in the locus without any 3D chromosome conformation information. Moreover, CMs capture different CREs that contribute to expression of the same gene in a cell type-specific manner. A notable example comprises the *ICOSLG* locus, where a putative enhancer 50 kb upstream of *ICOSLG,* previously shown to interact in 3D with the *ICOSLG* promoter [43], is associated with *ICOSLG* expression in Monocytes and Neutrophils. This CRE does not contribute to gene expression in T cells, where expression is instead most associated with signal exclusively around the *ICOSLG* promoter (**Fig 3e**). This is also reflected in the localization of the cmQTL in these respective cell type-specific CREs, indicating how cmQTLs allow to unravel cell type-specific regulation of gene expression. Finally, it is also conceivable that the same CRE drives expression of a gene, but that a cell type-specific element assists in this process. A notable example is the *CD40* gene expressed in both Monocytes and Fibroblasts, where a monocyte-specific CM captures an additional putative enhancer element, and both elements are only impacted by a variant in Monocytes (**Fig S3.2j**). This shows that CMs can capture CREs that are impacted by genetic variants located in a distal CRE in a cell type-specific manner.

Altogether, these analyses provide conceptual examples of how CM mapping aids in obtained a fine-grained understanding of the regulatory mechanisms underlying (cell type-dependent) gene expression.

### cmQTLs are associated with binding of cell type-specific TFs

Next, we assessed the cell type specificity of cmQTLs and observed that hQTLs were more frequently hQTLs in another cell type (or in LD (R2 > 0.8) with hQTLs) compared to cmQTLs. For example, 31.6% of H3K27ac hQTLs vs 19.4% of cmQTLs were shared between Monocytes and Neutrophils, indicating that cmQTLs capture more cell type-specific activity, as also revealed based on the enriched TFBSs (**Fig 2; Fig S4.1d**). As both hQTLs and cmQTLs can provide regulatory cues by disrupting or creating TFBSs, we compared histone QTLs that are also cmQTLs (which we will refer to as ‘hcmQTLs’ for simplicity) to QTLs that only impact a histone modification peak (hQTLs). Both QTL types displayed a similar genomic distribution (**Fig S4.1f**) yet hcmQTLs had on average a bigger impact on the peak height (i.e., higher beta values in a linear regression of genotype and normalized peak counts; **Fig S4.1g**). Both QTL types were enriched in cell type-specific open chromatin regions, with slightly higher specificity for hcmQTLs (**Fig S4.1h**), and hcmQTLs displayed stronger overlap (or in LD (R2 > 0.8) with cell type-specific expression QTLs (eQTLs; **Fig S4.1i**). Given that CM CREs enriched for TFBSs associated with cell type-specific TFs, we aimed to assess if hcmQTLs would also more often disrupt TFBSs for cell type-specific TFs compared to hQTLs. To do so, we compared the TF binding events that are disrupted by the QTLs using allele-specific binding (ASB) analysis [44](**Methods**). This revealed that a higher proportion of hcmQTLs is associated with ASB in any cell type compared to hQTLs with associations of QTLs with ASBs being partially cell type-specific. Notable examples include enrichments in ASB for the myeloid master regulator CEBPA/B in Monocytes and Neutrophils [29], for the mesoderm TF TCF21 in Fibroblasts [45] and the T(h1) and lymphoid factor TBX21 in T cells [46] (**Fig S4.2a**). In addition, many of the ASB events were more frequent in the hcmQTL group, indicating that hcmQTLs are associated with regions of dense(r) TF binding (**Fig S4.2b**). We complemented these analyses by assessing which TFs are more often binding (based on ChIP-seq data in any cell type) in a 200 bp window around the hcmQTL compared to hQTLs [47]. This validated the ASB observations in terms of binding of cell type-enriched TFs at hcmQTLs, such as IRF4 and BCL6 TFs in LCL, JUN/FOS in Fibroblasts and C/EBP in Monocytes and Neutrophils (**Fig S4.2c**). Together, these observations further indicate that CMs tend to arise in a cell type-specific manner driven by the binding of cell type-related TFs.

### Deconstructing regulatory hierarchies at autoimmune disease GWAS loci using CMs

Compared to QTLs exclusively impacting histone PTMs, hcmQTLs have a stronger impact on peak height, disrupt peaks that are most strongly associated with gene expression and are more likely to overlap cell type-specific eQTLs. We reasoned that hcmQTLs could thus be used to improve our understanding of disease predisposition. First, we computed the overlap (shared variants or in LD with R2 > 0.8) of hcmQTLs and hQTLs with variants reported in the GWAS Catalog [48] and compared the observed/expected ratio for the two types of QTLs with respect to overlap with the GWAS Catalog. This revealed several associations that occurred more frequently for hcmQTLs in the tested cell type, such as ‘Monocyte counts’ in Monocytes, ‘Lymphocyte counts’ in T cells and ‘Neutrophil percentage of white cells’ for Neutrophils (**Fig S4.3a**), further underlining the cell type specificity of cmQTLs. To assess how cmQTLs can assist in understanding regulatory logic at disease loci, we focused on autoimmune disease as these have their origin in immune cell types and a large number of GWAS summary statistics is available. Specifically, we used a list of 340 GWAS loci where co-localization of GWAS has been observed with at least an eQTL, hQTL, methylation QTL or a splicing QTL [49]. We then assessed co-localization of the cmQTLs on these loci with GWAS summary statistics for ankylosing spondylitis (AS), Celiac Disease (CEL), Crohn’s Disease (CD), Juvenile dermatomyositis (DM), Inflammatory Bowel Disease (IBD), Multiple Sclerosis (MS), primary biliary cirrhosis (PBC), psoriasis (PSO), Rheumatoid Arthritis (RA) Systemic Lupus Erythematosus (SLE), Type 1 Diabetes (T1D) and Ulcerative Colitis (UC) [50,51,60,52–59]. The majority of these loci (n=275, 80.9%) contained a CM in at least one cell type (**Fig 4a**). We applied a Bayesian co-localization approach [61] and defined co-localization as a Posterior Probability (PP) > 0.8 and at least one variant with a p-value < 1e-5 for both cmQTL (without distinguishing hQTLs and hcmQTLs) and GWAS association. This revealed co-localization of 59% (n=161) of cmQTLs with GWAS signal for at least one autoimmune disease (**Fig 4b; Fig S4.3b**), with more than half of the mapped CMs as well as cmQTL-GWAS co-localization being cell type-specific and the large majority (in LD with) an eQTL [26] in at least one tested cell type (**Fig 4c-e**). Next, we aggregated the co-localizations on a per-gene basis as one gene can be associated with multiple GWAS variants, resulting in 106 GWAS gene – cmQTL co-localizations (**Fig 4e**, some examples of co-localizations in **Fig 4f**). A notable example comprises the *SKAP2* locus where there are two associated candidate cmQTLs (rs2960785 and rs774267) with equal p-value of association with the aCM score, as well as expression of the CM-embedded genes *SKAP2* and *HOXA1* in Neutrophils (**Fig 4g-h**). We found that only rs2960785 localizes inside an intergenic enhancer region between *SKAP2* and *HOXA1*, indicating that rs2960785 is the likely causal variant. In addition, rs2960785 is highly blood-specific (**Fig S4.3c**), creates a putative GABPA binding site (CGGAAG)[62] (which is a direct interactor of the myeloid TF CEBPA [63]) and is associated with ASB of CEBPA in AML[44], providing direct insights into the molecular mechanisms that likely underlie how this genetic variant impacts the locus in a cell type-specific manner. For the *C3* and *TNFSF14* locus, the highest associated cmQTL is rs339392, which localizes in the *C3* promoter where it impacts chromatin activity at the locus, the expression of *C3* itself and to some extent also the expression of *TNFSF14* in LCLs (**Fig 4i-j**). The variant is also an eQTL for C3 in LCLs (**Fig S4.3c**) and is predicted to create a PU.1 / ETS binding motif (**Fig S4.3d**). The variant rs339392 was recently confirmed as a PU.1 binding QTL in LCL [64], and this impact on PU.1 binding extends to the other peaks embedded within the CM (**Fig 4i**), showing how a DNA base change can impact both histone modifications and TF binding on the focal CRE as well as associated distal CM-embedded CREs. A final example shown here involves the *TRIM14* locus, where the variant rs7867966 in the *TRIM14* promoter impacts the aCM score and the expression of *TRIM14* as well as the nearby gene *CORO2A*, likely owing to the impact of the variant on the promoter of *CORO2A* that is also embedded in the CM (**Fig 4k-l**). Similar to rs2960785, rs7867966 is an eQTL specific to blood cells (**Fig S4.3c**) and predicted to disrupt a C/EBP binding site (**Fig S4.3e**). Together, these example loci conceptually illustrate how mapping of CMs and associated QTLs can be used for the mechanistic interpretation of disease-associated non-coding variants, here exemplified by autoimmune disease-associated genetic variants. In addition, it allows to narrow down the number of candidate variants within a locus, as frequently more than one hQTL per locus can be identified (**Fig 4g, i, k**; red triangles).

**Figure 4.**
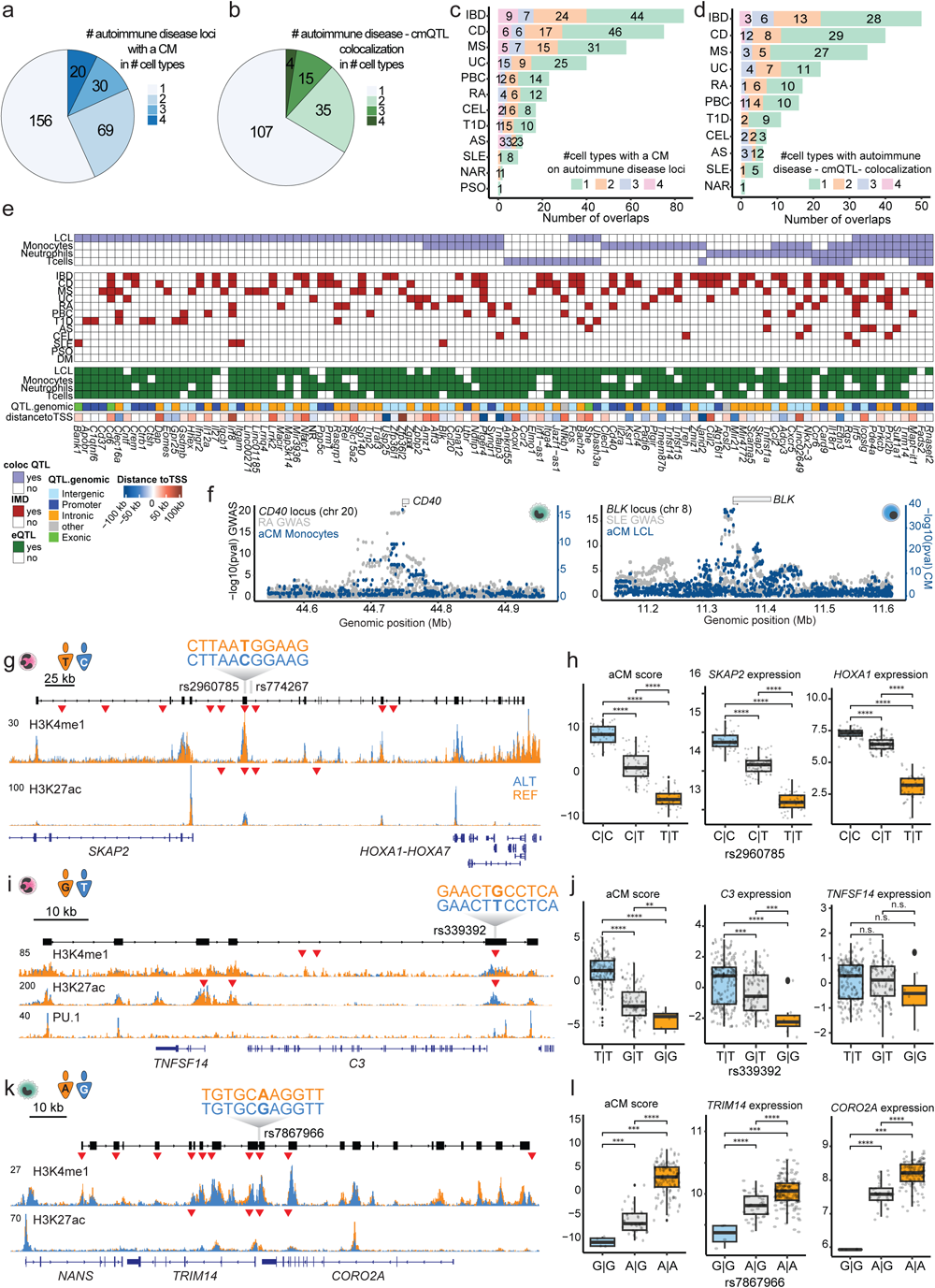
Mapping the regulatory logic at autoimmune disease risk loci using CMs. **a.** The number of autoimmune disease-associated GWAS loci where a CM was mapped in any of the four immune cell types. **b.** The number of loci where the cmQTL colocalized with a GWAS QTL for least one autoimmune disease (Posterior Probability (PP) > 0.8 and at least one variant that has a p-value of 1e-5 for both GWAS and variant-aCM association). **c.** The number of loci where a CM was mapped categorized per autoimmune disease. **d.** The number of loci with a cmQTL-GWAS colocalization categorized per autoimmune disease. **e.** Summary overview of autoimmune disease risk loci that harbour a CM with the associating cmQTL colocalizing with the GWAS variant for at least one disease. The color legend can be found at the lower left of the panel. Abbreviations are: spondylitis (AS), Celiac Disease (CEL), Crohn’s Disease (CD), Juvenile dermatomyositis (DM), Inflammatory Bowel Disease (IBD), Multiple Sclerosis (MS), primary biliary cirrhosis (PBC), psoriasis (PSO), Rheumatoid Arthritis (RA), Systemic Lupus Erythematosus (SLE), Type 1 Diabetes (T1D) and Ulcerative Colitis (UC). **f. *Left***: Example of colocalization of rheumatoid arthritis (RA) GWAS signal and the cmQTLs at the *CD40* locus in Monocytes. ***Right***: Example of colocalization of systemic lupus erythematosus (SLE) GWAS signal and cmQTLs at the *BLK* locus in LCLs. **g.** Example depicting the *SKAP2* locus in Neutrophils, one example individual per genotype. **h.** CM activity, *SKAP2* and *HOXA1* expression stratified by genotype of the highest-ranked candidate-associated variant rs2960785. **i.** Example depicting the *C3* locus in Neutrophils, one example individual per genotype. **j.** CM activity, *C3* and *TNFSF14* expression stratified by genotype of the highest-ranked candidate-associated variant rs339392. **k.** Example depicting the *TRIM14* locus in Monocytes, one example individual per genotype. **l.** CM activity, *TRIM14* and *CORO2A* expression stratified by genotype of the highest-ranked candidate-associated variant rs7867966. In **g**, **i** and **k,** the red triangles represent the localizations of histone QTLs. In the boxplots, each dot represents one individual and p-values were calculated using a Wilcoxon test. P-value indications are non-significant (ns) for p-value > 0.05, * for 0.01 < p-value ≤ 0.05, ** for 0.001 < p-value ≤ 0.01, *** for 0.0001 < p-value ≤ 0.001, **** p-value ≤ 0.0001.

### Chromatin modules reveal how genetic variants can impact expression of immune surface markers

Surface markers represent the molecules essential for immune cell interactions and consequently function, and several associations between non-coding variants and cell surface marker expression have already been reported [65]. Altered regulation of surface marker expression can affect how immune cells interact to combat infection and also impact the efficacy of treatment modalities that target these markers such as immunotherapy regimes. To assess the mechanisms underlying epigenomic regulation of surface marker expression, we aimed to investigate links between surface marker expression, CMs and associated QTLs. We used a list of 185 surface markers consisting of all Cluster of Differentiation (CD) and Tetraspanin molecules [66]. For 69 of these loci, we identified a CM in at least one cell type, and for 44 loci, a cmQTL was found in at least one cell type (**Fig S4.4a**). For example, for the CM spanning *CD207* in Monocytes, the top associated cmQTL rs11126300 localizes in an intergenic enhancer region between *CD207* and *CLEC4F*, where it modulates the epigenome of a range of putative CREs as well as the expression of both *CLEC4F* and *CD207* (**Fig S4.4b-d**). While CD207 is only expressed very lowly, *CLEC4F* is expressed higher and important for monocyte-derived Kupffer cells [67]. Another relevant example is the *CD93* locus in Neutrophils, where a CM spans a range of genes. The top ranked candidate associated variant rs844881 localizes in a peak at the 3’ UTR of the lncRNA *LINC00656*. This variant has a strong impact on the activity of the respective aCM, and also leads to strong reduction of the expression of *THBD*, *CD93* and *LINC00656*. Interestingly, we did not observe an expression change for *NXT1* and *NAPB*, which are adjacent genes that are not part of the respective CM (**Fig S4.4e-g**). The variant rs844881 shows a strong association with the development of varicose veins in the FinnGen cohort (p = 5.5e-5)[68]. This could be related to the impacted expression of the *THBD* gene, as altered *THBD* expression in Neutrophils has been implicated in the development of venous thrombosis [69] and thus together provides a putative mechanism of how rs844881 could impact the risk on varicose veins by impacting *THBD* expression in Neutrophils. Finally, the example of the *CD9* locus shows that even though the same genetic variant impacts an intergenic enhancer in different cell types, it is only when adjacent CREs are included in the CM that this results in an impact on gene expression (**Fig S4.4h-i**).

Altogether, these analyses reveal how genetic variants can impact CMs around surface markers and potentially result in functional variation in immune cell function with impact on disease susceptibility.

### Context-dependent formation of CMs in chronic lymphocytic leukaemia

Recent works revealed that non-coding genetic variants can impact the prognosis of chronic (CLL) and acute (ALL) lymphocytic or myeloid (AML) leukaemia [70–72]. For example, a variant in the *AXIN2* locus was shown to activate multiple enhancers (within a CM) as well as the expression of *AXIN2* which was linked to better survival rates of CLL patients [21]. Another study on ALL reported a variant in a *GATA3* enhancer that impacts multiple regions around GATA3, resulting in higher *GATA3* expression in *cis* and downstream 3D genome rearrangements [72]. As such, we aimed to assess if CMs could capture CLL-specific-induced epigenome configurations. For this purpose, we uniformly analysed epigenome data from LCLs (immortalized B cells in a non-disease context, n=317 individuals) and CLL (immortalized B cells in the context of cancer, n=105 individuals) (**Fig 5a**). Similar to the strategies applied for the five cell types (**Fig 2a**), we defined a common set of peaks based on H3K27ac enrichment in both contexts and used these for CM mapping. Since CMs capture variable signal, we reasoned that calculating the variance of each CM in both LCL and CLL in combination with the interindividual variability in gene expression would allow identification of genomic loci in which both the local chromatin environment as well as expression of the embedded genes are distinct between LCL and CLL (**Fig 5b; Fig S5a-b**). Such loci are of special interest since they may contribute to the leukemic state of these cells, and some of the CLL-specific locus-gene combinations identified in these analyses include previously implicated genes such as *CTLA4* and *WNT3* [73,74] (**Fig 5b**). The locus-gene combination that we found is most specific to CLL is the *OSBPL5* locus (**Fig 5b**). While not active in LCL, we observed that the *OSBPL5* promoter as well as several flanking enhancers are activated in a subset of CLL patients (**Fig 5c**). By associating this region to the underlying genotype (n=35 individuals for which both H3K27ac ChIP-seq and genotype data are available[21]), we identified a common candidate non-coding germline variant (rs895555, MAF=45%) in an *OSBPL5* intron that is strongly associated with the activation of the locus (**Fig 5d-e**). The T allele of rs895555 creates the core of a putative Forkhead (FOX) TF binding site ((<u>T</u>/C)GTTT) (**Fig S5c**), which is of interest given the implication of this TF family in the development and progression of B cell malignancies [75]. The expression of *OSBPL5* is a prognostic marker for overall survival in the context of mutated IGHV status (**Fig 5f**), and has also recently been identified as the strongest predictive gene expression marker for time to progression after chemoimmunotherapy treatment of CLL [76]. Thus, our findings indicate how a genetic variant might impact disease progression in a personalized manner through induction of a CM at the *OSBPL5* locus. The second-highest variable gene-locus combination was the *COBLL1* locus (**Fig S5d-f**), for which the strongest associated variant is a common (MAF = 17%) intronic TATA duplication in a subset of CLL individuals. *COBLL1* expression is associated with survival in chronic lymphocytic leukaemia (**Fig 5g**) and has also been linked to survival probability in several other cancer types including chronic myeloid leukaemia [77]. Finally, we analysed TF binding profiles in the CREs included in CLL or LCL CMs, compared to simulated reference CMs. We observed enrichment of many similar types of TFs, related to B cell function such as IRF, RUNX and EBF1 (**Fig S5g-h**), but also specific enrichment of factors such as FOXO1 (as also seen for rs895555) in CLL which have been implicated in CLL and harbour driver mutations [75]. Despite the limited number of genotypes available for CLL, we provide evidence that CMs can be used to assist in understanding how non-coding variants can impact local epigenome regulation related to disease-relevant genes.

**Figure 5.**
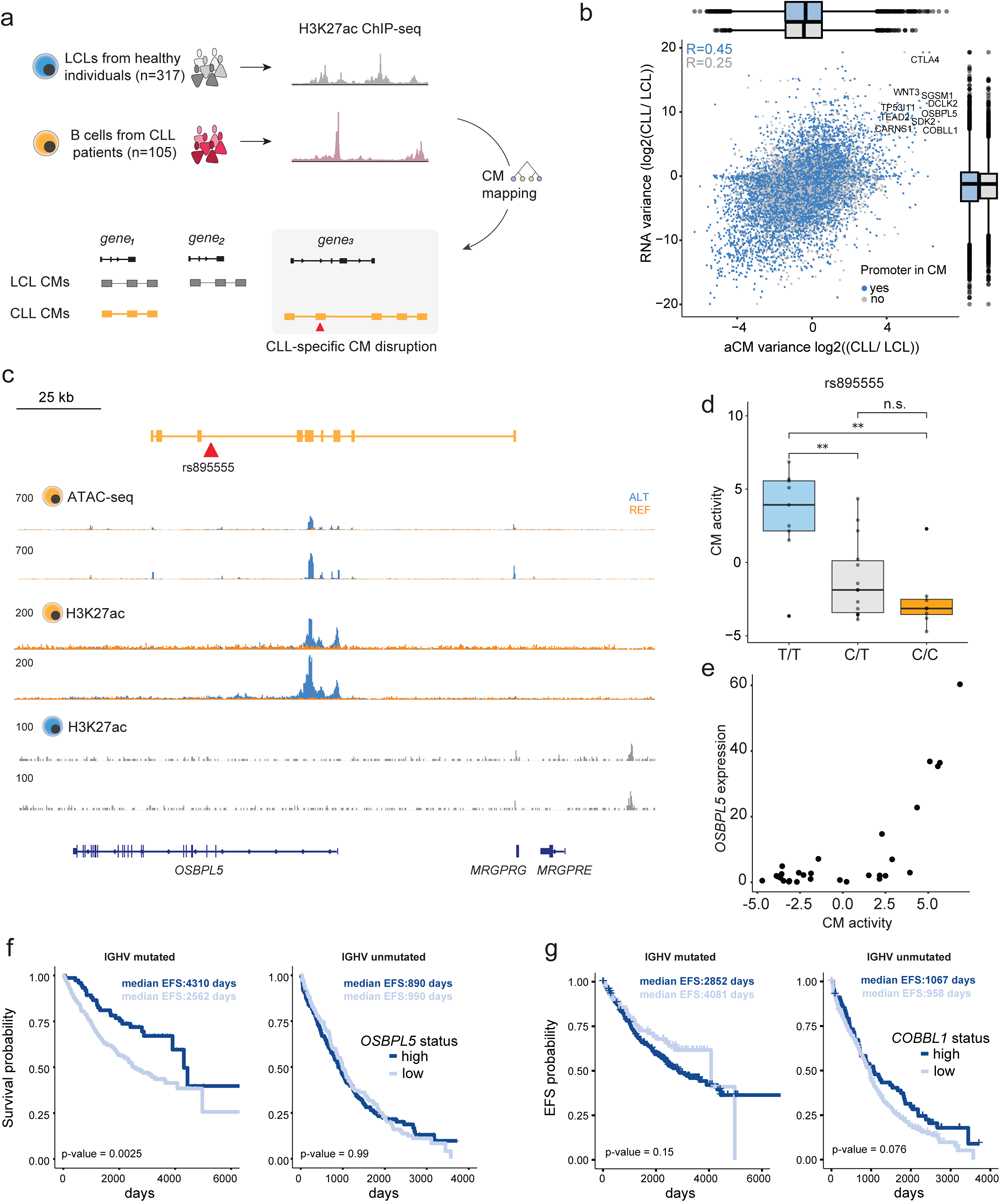
Activation of genomic loci in a subset of CLL patients. **a.** Strategy for mapping CMs using available ChIP-seq for H3K27ac in LCL (n=317) and CLL (n=105) samples. **b.** Log2 ratio of interindividual variance of aCM activity (x-axis) and interindividual variance in expression of the gene(s) embedded within the CM. Colors represent whether the promoter of the gene is part of the CM. Correlation values are Pearson correlation. Boxplots represent the distribution of the interindividual variance. **c.** Example depicting the *OSBPL5* locus whose embedded gene’s expression is induced in a subset of CLL patients. **d.** CM activity stratified by genotype of the highest-ranked candidate-associated variant rs895555. **e.** Correlation between aCM activity and *OSBPL5* expression. **f-g.** Event-free survival of CLL patients stratified on *OSBPL5* (**f**) or *COBLL1* (**g**) expression for both IGHV-mutated and -unmutated CLL status. P-values were obtained using a log-rank test. P-value indications are non-significant (ns) for p-value > 0.05, * for 0.01 < p-value ≤ 0.05, ** for 0.001 < p-value ≤ 0.01.

## Discussion

In this work, we established a set of computational guidelines to map CMs and harmonize the output formats across methods, which makes CM mapping accessible to the community (see **data availability**). We used this to comprehensively assess regulatory coordination in six cell types and hundreds of individuals in a genome-wide manner, which revealed extensive cell type specificity of regulatory variation. We provide mapped CMs in an accessible format (see **data availability**) and show how these can be used to disentangle the different (epi)genomic modalities, starting from genetic variants to TFBSs, histone modifications on local and distal CREs and finally gene expression. Our results substantiate the hypothesis of cooperation between local and distal CREs within TADs, which we observed using both epigenome data and enrichment in 3D interactions. At the sub-TAD scale, we argue that mapping of CMs thus provides an approach complementary to 3C-based applications for determining CRE interactions within TADs in a genome-wide manner [17–20]. An advantage of mapping CMs is that CREs are grouped into functional units, and that CMs therefore allow to provide mechanistic rationales regarding how non-coding variants can impact the epigenome of local and distal CREs as well as the expression of genes embedded within the respective CM. We provide examples of such analyses in the case of GWAS loci for autoimmune disease, surface marker expression variation on immune cells and identification of candidate variants driving prognostic marker expression in CLL patients.

Ultra-resolution 3C-based assays or microscopy strategies have revealed in specific loci that the removal of the canonical proteins involved in regulating 3D genome architecture, such as CTCF and Cohesin, has limited impact on the establishment of interactions between CREs [78–83]. Moreover, structural proteins such as CTCF are ubiquitously expressed, and tend thus on themselves to be unable to define the cell type-specific CRE cooperativity within CMs. Together, this raises questions of how regulatory interactions within a TAD environment are established in a cell type-specific context. An attractive hypothesis is that this is mediated by cooperating TFs, which often are expressed in a cell type-or lineage-specific manner [84], followed by TF-mediated recruitment of general co-activators and chromatin modifiers [17]. The observations on the cell types assayed here suggest that CMs are established by classes of more ubiquitously expressed TFs that bind CM-embedded CREs that are shared across multiple cell types (here defined as anchors) in combination with cell type-specific TFs that bind CM-embedded CREs whose activity is restricted to one of just a few cell types. Such cooperative binding of cell type-/lineage-specific and core CM TFs [33] that reflect a specific cell state may thus mediate CRE interactions within and beyond the 3D genome organization established by CTCF and Cohesin. This observation is also consistent with the notion that cmQTLs frequently disrupt the binding sites of cell-type specific TFs, which then could impact histone modification deposition and cooperation between CREs within a CM in a cell type-specific manner. This finding complements recent observations showing that genetic variants impact CRE interactions relevant for establishing gene expression through direct perturbation of TF binding to enhancers [85] and that TFs are the main drivers of gene expression cooperativity [86].

## Conclusions

Altogether, we demonstrate that CMs provide a means to shed light on the mechanisms underlying gene expression and gene regulatory variation with high-throughput epigenome data in a range of genetic backgrounds, provided the availability of a sufficient number of assayed samples. Future applications of CM mapping could include defining CM plasticity within closely related cell types during cellular differentiation, or rather in different species to assess how conserved local genome organization is. With the constantly decreasing cost of high-throughput sequencing and easy-to-implement epigenome tools, we advocate for the value of CM mapping in future studies where epigenome data is being generated in large sample cohorts as part of the overarching goal to understanding how regulatory variation contributes to complex traits and disease [18,19,26,65,87]. We believe that the guidelines, executable code and interpretable output provided here will highly facilitate this process

## Methods

### CM mapping

The methods used in this work required adjustments (or full implementation, as in the case of VCMtools). The CM mapping approach described in Waszak et al. [18] was implemented in python (hereafter referred to as VCMtools) and adapted to account for the cell type-specific background by using empirical p-value correction followed by FDR thresholding. Namely, for each chromosome, for two peaks, whose centers are located at the max distances of 500 Mb from each other, we calculated and stored correlation values of the respective peak height profiles across individuals. Then, we calculated the average background correlation (*μ*_*bg*_) and its standard deviation (*ς*_*bg*_) for all peak pairs across all chromosomes. This allowed us to calculate the empirical p-value = 1 – CDF (correlation; μ_*bg*_, σ_*bg*_) (scipy.stats.norm.cdf(correlation, loc=*μ*_*bg*_, scale=*ς*_*bg*_)), which we further corrected with the Benjamini-Hochberg procedure. CMs were identified by selecting isolated components after filtering peak-to-peak correlations based on the specified corrected p-value threshold (p-value <= 0.001). Clomics did not require major modifications, except for the conversion of the output tree file into a BED file format. By following the described methods in the supplementary section of Kumasaka et al., 2019 [20], we were able to implement some of the key methods included in the PHM package (https://github.com/natsuhiko/PHM/), including the DAG construction for inference of hierarchies between peaks categorized into the causality hypothesis (posterior probability of causal interaction >= 0.8). Data preprocessing and CM mapping scripts are available via GitHub (https://github.com/DeplanckeLab/Chromatin_modules).

### Randomization strategy for investigating the number of samples required to map CMs

The robustness of the CM mapping methods was evaluated with respect to the number of individuals in every cell type. We chose the smallest chromosome chr22 and ran each of the methods on a subset of samples of size *s* ∈ {*x* × 25 | *x* = 1, 2, …, 12}, where each of the subsets was randomly sampled five times, resulting in 12 x 5 runs per method. For each execution run, we measured the number of CMs, elapsed time, memory consumption, median module length and percentage of CMs with a cmQTL.

### Reproducibility analysis of mapped modules across cell types and methods

To compare CMs, we developed the approach based on the F1 score[88]. We started by calculating the Jaccard overlap between the peaks at the level of base pairs. Then, for each pair of CMs 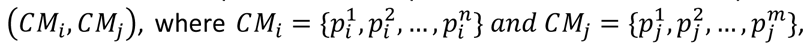, we we created a matrix of size *N* × *M*, where *N* = |*CM*_*i*_| is the number of peaks in *CM*_*i*_ and *M* = |*CM*_*i*_| is the number of peaks in *CM*_*i*_. If there is an overlap between a *p*^*k*^_i_ and *p*^l^_j_, then the value in the *k*-th row and l-th column of the *i* *i* matrix *J*, is defined as a Jaccard index *J_k,l_* 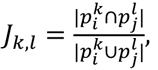, where *p*^*k*^_*j*_ and *p^l^_j_* are defined as sets of unique genomic positions.

To obtain a similarity/reproducibility score for a pair of CMs, we first calculated the average of maximum values along the rows (1) and columns (2) of matrix J as follows:

(1)

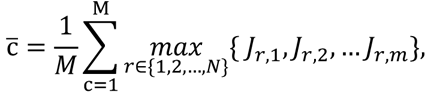
(2)

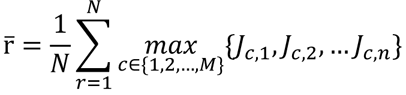

Finally, the similarity/reproducibility score was defined as 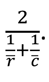

### CM overlap with A/B compartments, TADs, chromatin state annotations and genes

TAD and A/B compartment coordinates (for GM12878, GSE63525) were overlapped with CM coordinates using *bedtools* [89] function *intersect* with parameter *-f 1.0* to report only those CMs that were fully located in TADs or A/B compartments. CRE annotations for LCLs (GM12878) were obtained from ChromHMM [10] and SCREEN [90] databases. Annotated CRE coordinates were overlapped with CM and non-CM peaks with *bedtools* without restrictions on the overlap size. Then, per each CRE annotation category and per peak, we calculated the fraction of peak length overlapping with the respective annotated CRE. For each CRE category, we performed Fisher’s exact test on the contingency table constructed for CM peaks and non-CM peaks in a category of interest (non-significant (ns) for p-value > 0.05, * for 0.01 < p-value ≤ 0.05, ** for 0.001 < p-value ≤ 0.01, *** for 0.0001 < p-value ≤ 0.001, **** p-value ≤ 0.0001). Gene coordinates were downloaded from the GRCh37 (hg19) assembly. For every gene, we defined the promoter region as +-500bp from the TSS, and gene body as gene coordinates without promoter region. Gene coordinates (promoter + gene body) were overlapped with CM peaks with *bedtools* without restrictions on the overlap size.

### Simulation strategy for CMs

To move away from a single or paired element-defined background, we devised a simulation strategy to construct a background set of CMs resembling mapped regions of covariable CREs in terms of their intrinsic properties, which include:

1. Number of regulatory elements in a CM,
2. Length of a CM,
3. Total base pair length of elements in a CM,
4. Overall peak signal variability in a CM,
5. H3K27ac peak fraction,
6. CM location in A or B compartments and TADs (Optional).

Prior to simulating the background set of CMs, we define the reference set of CMs as the set of all CMs if A/B compartment and TAD data (pt.6) is not available. For the consistency of the analysis described in the paper, we did not use the A/B compartment and TAD constraint since the data was not available for all cell types. In the case when this data is available, the reference set of CMs is defined by overlapping the mapped CMs with TAD boundaries and stratifying the CMs based on their localization in A/B compartments. This allows to narrow down the set of CMs to putative functional regions by conditioning on TADs, and account for potential structural differences of CMs located in A or B compartments.

Every CM in the reference set is characterized by a vector of features described above in points 1-6. Then, for every CM from the reference set of size n, we randomly sampled a peak from the non-CM peak set, and n-1 peaks in the peak neighborhood. In the case of simulation of CMs for the PHM output, we started by randomly sampling a peak classified into one of the single QTL hypotheses. The number of peaks per histone mark in the set of sampled non-CM peaks was perfectly matched with the respective reference CM histone mark peak frequencies. We repeated the procedure 10 times per CM, resulting in 10 sets of sampled peaks per CM, where each set represents a candidate simulated CM. These candidates for simulated CMs were characterized with the same feature set as for CMs. For every CM in the reference set, we searched for the five most similar simulated CMs among all generated candidates of the same size (same number of peaks), with respect to the feature vector, by applying Approximate Nearest Neighbors search (Annoy package https://github.com/spotify/annoy). After matching CMs with simulated CMs, we filtered out simulated CMs with more than 10% variation in GC content, CM length, total base pair length of peaks as compared to the reference CM. This allowed us to obtain the final set of paired mapped and simulated CMs.

### Quantification of peak interactions in Hi-C and Micro-C data

Micro-C data was aligned using BWA v0.7.17 [91]. Ligation events were determined using pairtools parse with parameters --min-mapq 40 --walks-policy 5unique --max-inter-align-gap 30. PCR duplicates were removed using pairtools dedup followed by the generation of .bam and .pairs files. The pairs files were used to generate cooler files using cooler cload pairix using default parameters [92]. For the constructed set of paired mapped and simulated CMs, interaction frequencies for peaks in CMs were quantified in Hi-C and Micro-C data at 500bp, 1kb and 5kb resolutions by first fetching the bins overlapping individual peaks and then averaging the signal within the bin overlap area for all possible peak pairs in CM.

### Differential TFBS enrichment

We used command line UniBind TFBS differential enrichment tool (*UniBind_enrich.sh* script with *twoSets* parameter and *hg38_robust_UniBind_LOLA.RDS* motif set). Prior to running the script, peak files were lifted from *hg19* to *hg38*. Depending on the task, the background set was changed, e.g., for differential TFBS enrichment in actual CM peaks vs simulated CM peaks, we used simulated CM peaks as a background set. For TFBS enrichments at CM peaks in cell type A vs cell type B, CM peaks from the cell type B were used as a background.

### ChIP-seq and ATAC*-*seq data processing

Raw fastq files were aligned to the hg19 genome using Bowtie2 v2.4.4 [93]. Unpaired reads were removed using samtools [94] and duplicates were marked and removed using picard v2.17.8 (http://broadinstitute.github.io/picard/). To create a common set of peaks per cell type, first a ‘meta’ bam file was created per epigenome assay (H3K4me1, H3K27ac or ATAC) by merging down-sampled (5 million reads per sample) bam files for each individual using samtools merge. The resulting meta bam files of the different cell types were then merged per epigenome assay resulting in the creation of the final merged bam files. Peak calling was performed on these files using MACS2 [95] using parameters ‘--broad’ -f BAMPE and a q-value cutoff of 0.01. The obtained universal peak set was used to count the number of reads for the respective epigenome assays per individual per cell type using FastReadCounter (https://github.com/DeplanckeLab/FastReadCounter). The resulting non-normalized count matrices were RPKM normalized using the rpkm function from edgeR v3.36.0 [96]. For data standardization, we used linear regression to remove the known covariates sex and age. As the LCL cohort consists of a merge of cohorts from individuals from different ancestries, the first 3 principal components of the genotypes (obtained using QTLtools pca [97]) were removed as well. To account for effects of unknown covariates, principal component analyses were performed on the RPKM-normalized count matrices and the first 10 PCs (in case of Monocytes, Neutrophils, Tcells and CLL) or the first 20 PCs (in case of LCL and FIB) were also removed during the regression. The residuals from the linear regression were kept and normalized using quantile normalization with the qqnorm function from the stats base R package. The resulting matrices were used as input for mapping of QTLs and CMs. RNA-seq files for LCL and FIB were processed in the same way, with variance stabilization using the DESeq2 R package (v 1.40.1) [98] instead of rpkm normalization.

### Calculation of the CM activity score (aCM)

For each CM, the embedded peaks and associated counts per individual were extracted to create a separate peak by individual matrix, followed by principal component analysis. Since the embedded peaks are correlated by definition, most variation in signal between the individuals is captured in the first principal component, and thus the values for this component were extracted and used as the aCM score.

### Mapping of quantitative trait loci (QTLs)

Mapping of QTLs was done using QTLtools [97] using the options QTLtools *cis* with the options -- permute 1000 and --normal. To account for the fact that genotype-phenotype associations are assayed for a large number of loci, the resulting FDR-adjusted p-values were further corrected using q-value correction using the qvalue R package (v2.24.0). Variants with a fdr-q-value corrected p-value under 0.05 were considered significant.

### Intersection of QTLs with GWAS

To allow for a complete overlap, the variants in linkage disequilibrium (LD) with the QTLs were determined using plink2 [99] based on the 1000G genotype file for individuals with a European ancestry. All variants within a 1 Mb window were considered and those with R2 > 0.8 retained as LD variants. The GWAS catalog for hg19 with 392271 associations (release 2022-05-11) was downloaded from the UCSC table browser and only variants that were present in the genotype file and the GWAS catalog were considered for downstream analyses. To compare histone QTLs that were also cmQTLs with hQTLs, the hQTL files were split into variants that were also cmQTLs themselves or in LD (R2 > 0.8) with cmQTLs (from then on referred to as hcmQTLs) and only hQTLs. Then, variants were overlapped with the GWAS variant file, and considered overlapping based on a direct match or LD (R2 > 0.8) with a GWAS variant. This yields the quantification of how many overlaps there are of the tested QTL set with each GWAS variant. By comparing this with the QTL set size and the total occurrence in the GWAS file, this allows to obtain the observed / expected ratio for each QTL set individually. A Fisher’s exact test was used to compare this ratio between hcmQTL and hQTL sets to identify the enriched terms. For colocalization of cmQTLs with GWAS variants, each variant in a 500 kb window around the CM was tested for the association with the aCM score. The resulting variants were matched with those present in the summary statistics for each of the tested autoimmune disease-associated variants. Colocalization was performed using Coloc v5.2.2 [61]. Variants with a posterior probability (PP4) > 0.8 and a p-value under 1e-5 for both the cmQTL and GWAS associations were considered as colocalizing.

### Intersection of QTLs with TF binding

QTLs were divided in hcmQTLs and hQTLs and 200 bp windows were created around each variant. The R package ReMapEnrich v0.99.0 [47] was used to identify TFs that were found to bind in these windows using ChIP-seq in any human cell type. Enrichment was assessed using the hcmQTL set as input with the hQTL-only set as reference, thus serving as reference regions. Allele specific binding (ASB) events for 1073 TFs were downloaded from the Adastra database (release June 2022) [44]. For each TF, only ASB events with a reported FDR < 0.05 were retained. Identified hQTLs were stratified in hcmQTLs and hQTLs and intersected with the ASB events to identify the total number of ASB events for each TF for the two QTL groups. The number of ASBs for each TF was compared between the QTL groups using a Fisher’s exact test.

### Enrichment of QTLs in open chromatin regions

QTLs were divided in hcmQTLs and hQTLs and analyzed using Forge2 [100] via the associated web server (https://forge2.altiusinstitute.org/) using default parameters. Consolidated Roadmap Epigenome DNAseI hypersensitive sites were used for the enrichment analysis.

## Declarations

### Availability of data and materials

Raw fastq files for LCL and Fibroblasts were downloaded from the ArrayExpress Archive at EMBL-EBI (www.ebi.ac.uk/arrayexpress) (E-MTAB-3656 and E-MTAB-3657) and the European Genome-phenome Archive (European Genome-phenome Archive; https://ega-archive.org/datasets/). Monocytes, Neutrophils and T cell ChIP-seq fastq (EGAD00001002670, EGAD00001002672, EGAD00001002673) and genotype data (EGAD00001002663) were obtained from the European Genome Archive. ChIP-seq and ATAC-seq data for CLL were downloaded from the European Genome Archive (EGAD00001004046). Processed RNA-seq files for Monocytes, Neutrophils and T cells were downloaded from ftp://ftp.ebi.ac.uk/pub/databases/blueprint/blueprint_Epivar/ [26]. Processed RNA-seq files for CLL were downloaded from http://resources.idibaps.org/paper/the-reference-epigenome-and-regulatory-chromatin-landscape-of-chronic-lymphocytic-leukemia [87]. Genotypes from the 1000 genomes were obtained from https://www.internationalgenome.org/data. All autoimmune disease summary statistics were obtained from the GWAS catalog (https://www.ebi.ac.uk/gwas/). Hi-C contact matrices, A/B compartments and TADs for GM12878 were downloaded from GEO (GSE63525) [3]. Processed ultra-high resolution Hi-C data for GM12878 at 500bp was obtained from ENCODE [101]. Micro-C data was obtained from Dovetail Genomics (https://micro-c.readthedocs.io/en/latest/data_sets.html). ChIP-seq tracks for TFs in the bigwig format for signal p-value of merged replicates were downloaded from ENCODE for GM12878 and IMR-90 for the following TFs: EBF1 (ENCFF248XJC for GM12878), PAX5 (ENCFF759XQV for GM12878), BHLHE40 (ENCFF429WGS for IMR-90, ENCFF816QSI for GM12878), CTCF (ENCFF583IZF for IMR-90, ENCFF749HDD for GM12878). All the code for mapping CMs using VCMtools, Clomics or PHM is publicly available at GitHub (https://github.com/DeplanckeLab/Chromatin_modules).

## Acknowledgements

We want to thank the data generators from the following entities: The International Cancer Genome Consortium (ICGC), The Gencord cohort, The Blueprint Consortium and The Genotype-Tissue Expression (GTEx) Consortium. We further want to acknowledge the participants and investigators of the FinnGen study. This work was supported by a Swiss National Science Foundation grant (no. 310030_197082) to B.D. and by Marie Sklodowska-Curie fellowships for O.P. (no. 860002), G.v.M. (no. 101026623), J.F.K. (no. 895426) and W.S. (no. 101028476). The authors thank all the Deplancke’s lab members for valuable discussions.

