## Supplementary_Figures for "Non-coding variants impact *cis*-regulatory coordination in a cell type-specific manner"

**Supplementary Material**

Supplementary Figures. **Figure S1.1 – Figure S5.**

Supplementary Table 1. **Chromatin modules mapped in LCLs using VCMtools, Clomics and PHM.**

Supplementary Table 2. **Chromatin modules mapped in five cell types using Clomics.**

Supplementary Table 3. **cmQTLs for chromatin modules in five cell types.**

Supplementary Table 4. **Categorization of TFBSs in CM CREs.**

Figure S1.1

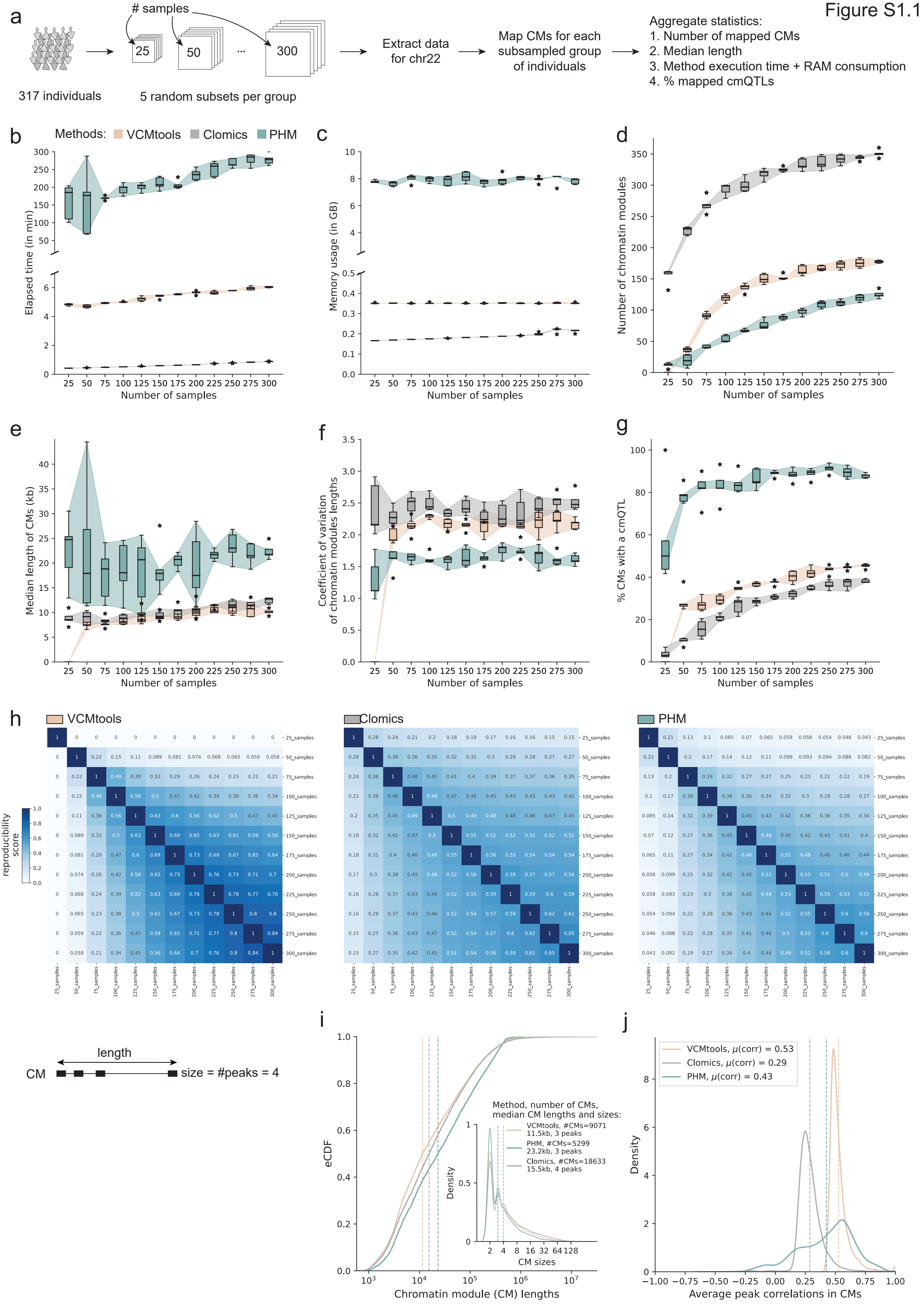

a
### samples
25
50
...
300
317 individuals
5 random subsets per group
Extract data for chr22
Map CMs for each subsampled group of individuals
Aggregate statistics:
1. Number of mapped CMs
2. Median length
3. Method execution time + RAM consumption
4. % mapped cmQTLs
b
Methods: VCMtools Clomics PHM
Elapsed time (in min)
Number of samples
c
Memory usage (in GB)
Number of samples
d
Number of chromatin modules
Number of samples
e
Median length of CMs (kb)
Number of samples
f
Coefficient of variation of chromatin modules lengths
Number of samples
g
% CMs with a cmQTL
Number of samples
h
VCMtools
Clomics
PHM
reproducibility score
i
eCDF
Chromatin module (CM) lengths
Method, number of CMs, median CM lengths and sizes:
VCMtools, #CMs=9071 11.5kb, 3 peaks
PHM, #CMs=5299 23.2kb, 3 peaks
Clomics, #CMs=18633 15.5kb, 4 peaks
Density
CM sizes
length
CM
size = #peaks = 4
j
VCMtools, μ(corr) = 0.53
Clomics, μ(corr) = 0.29
PHM, μ(corr) = 0.43
Density
Average peak correlations in CMs

**Figure S1.1. Performance comparison of CM mapping strategies and general CM characterization. a.** Schematic representation of the subsampling strategy on chr22 for evaluating the effect of the number of samples on different parameters such as the **b.** elapsed time of CM mapping (excluding data preprocessing time; no parallelization), **c.** maximum RAM occupancy (in Gb), **d.** number of mapped CMs, **e.** median CM length, **f.** coefficient of variation of CM length, **g.** percentage of CMs with chromatin module (cm)QTLs. **h.** Heatmaps of average reproducibility scores (F1-based), across five random groups per sample size batch, for CMs mapped with (***from left to right***) VCMtools, Clomics and PHM. **i.** Empirical Cumulative Density Function (eCDF) of CM lengths and sizes (***inset panels***) across methods. **J.** Distribution of average correlation of peaks within CMs across methods.

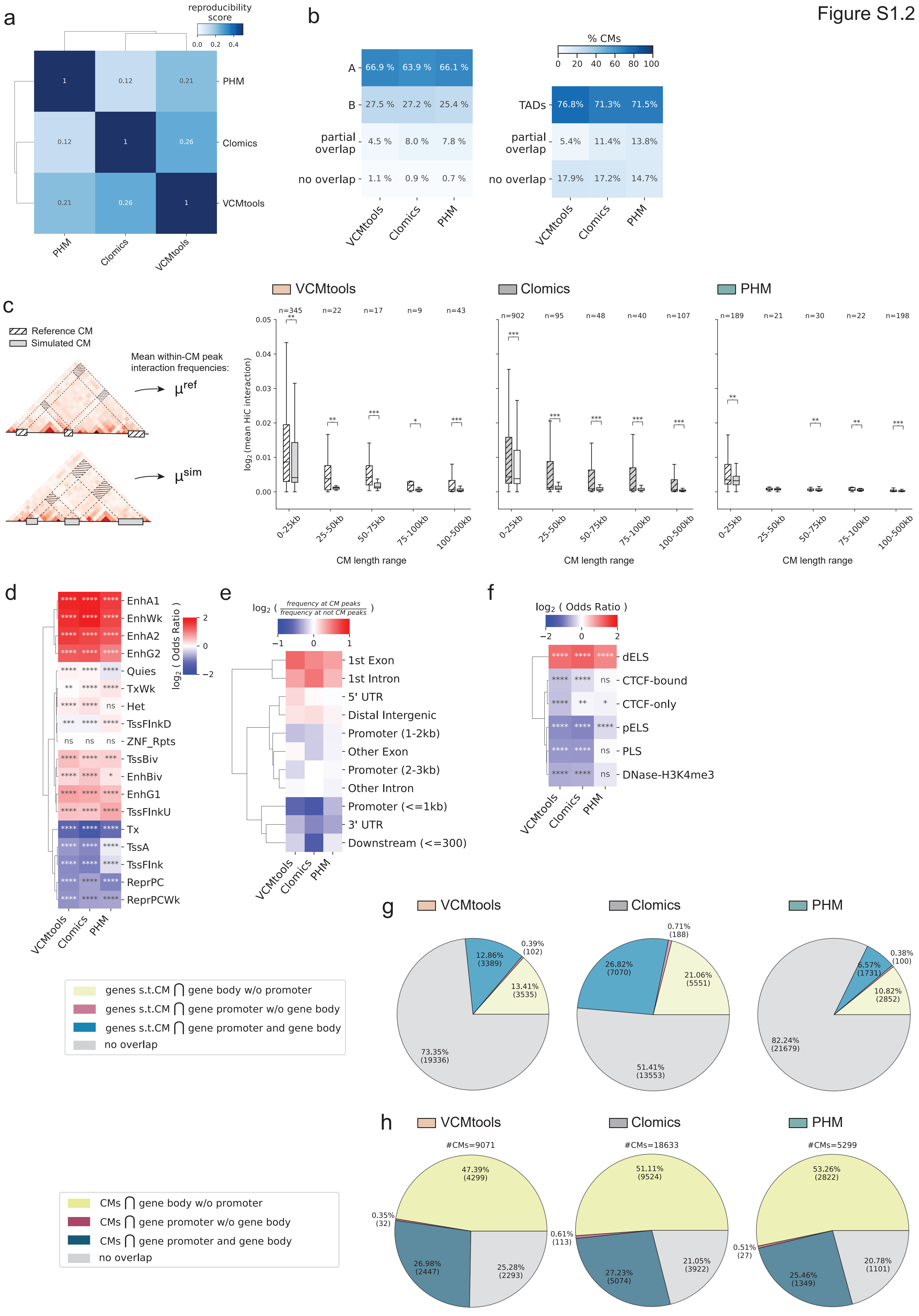
Figure S1.2
a
reproducibility score
0.0 0.2 0.4
PHM
Clomics
VCMtools
b
% CMs
0 20 40 60 80 100
A 66.9 % 63.9 % 66.1 %
B 27.5 % 27.2 % 25.4 %
partial overlap 4.5 % 8.0 % 7.8 %
no overlap 1.1 % 0.9 % 0.7 %
TADs 76.8% 71.3% 71.5%
partial overlap 5.4% 11.4% 13.8%
no overlap 17.9% 17.2% 14.7%
c
Reference CM
Simulated CM
Mean within-CM peak interaction frequencies:
μref
μsim
VCMtools
Clomics
PHM
log2 (mean HiC interaction)
CM length range
0-25kb 25-50kb 50-75kb 75-100kb 100-500kb
d
log2 ( Odds Ratio )
EnhA1
EnhWk
EnhA2
EnhG2
Quies
TxWk
Het
TssFlnkD
ZNF_Rpts
TssBiv
EnhBiv
EnhG1
TssFlnkU
Tx
TssA
TssFlnk
ReprPC
ReprPCWk
e
log2 ( frequency at CM peaks / frequency at not CM peaks )
1st Exon
1st Intron
5' UTR
Distal Intergenic
Promoter (1-2kb)
Other Exon
Promoter (2-3kb)
Other Intron
Promoter (<=1kb)
3' UTR
Downstream (<=300)
f
log2 ( Odds Ratio )
dELS
CTCF-bound
CTCF-only
pELS
PLS
DNase-H3K4me3
g
genes s.t.CM ⋂ gene body w/o promoter
genes s.t.CM ⋂ gene promoter w/o gene body
genes s.t.CM ⋂ gene promoter and gene body
no overlap
VCMtools: 12.86% (3389), 0.39% (102), 13.41% (3535), 73.35% (19336)
Clomics: 0.71% (188), 26.82% (7070), 21.06% (5551), 51.41% (13553)
PHM: 0.38% (100), 6.57% (1731), 10.82% (2852), 82.24% (21679)
h
CMs ⋂ gene body w/o promoter
CMs ⋂ gene promoter w/o gene body
CMs ⋂ gene promoter and gene body
no overlap
VCMtools #CMs=9071: 47.39% (4299), 0.35% (32), 26.98% (2447), 25.28% (2293)
Clomics #CMs=18633: 51.11% (9524), 0.61% (113), 27.23% (5074), 21.05% (3922)
PHM #CMs=5299: 53.26% (2822), 0.51% (27), 25.46% (1349), 20.78% (1101)

**Figure S1.2. CM mapping method comparison with respect to genomic CM localization, 3D interactions, CRE annotations and gene overlaps. A.** Clustered heatmap of average similarity scores between CMs mapped with different methods. The largest similarity score of 0.26 is between correlation-based approaches. The relatively low average score is a result of the difference in number of identified CMs that are considered in each pairwise comparison (cf. **Fig 1c, S1.1i**). **b.** Overlap of CMs with A/B compartments or TADs across methods. **c.** ***From left to right***: VCMtools, Clomics, PHM. Quantification of 3D interactions between CM peaks of mapped and simulated CMs with Hi-C data at 500bp resolution for CMs split by length. Stars indicate the respective adjusted p-value strength for the Wilcoxon test. The numbers indicate the number of reference and matched simulated CMs included in each category. **d.** Enrichment of CM peaks versus non-CM peaks in ChromHMM-annotated regions across methods. Color intensity corresponds to the log2 Odds Ratio, stars indicate the respective p-value strength for the Fisher exact test. **e.** Log2 frequency of CM peaks found in either of the ChIPseeker categories to the frequency of non-CM peaks found in the same category across methods. **f.** Enrichment of CM peaks versus non-CM peaks in SCREEN annotations of CREs across methods. **g.** ***From left to right***: VCMtools, Clomics, PHM. Percentages of genes overlapped by CMs (relative to all coding genes) falling into each category per method. **h.** ***From left to right***: VCMtools, Clomics, PHM. Percentages of CMs that cover total, no, or parts of genes, separated per method. P-value indications are non-significant (ns) for p-value > 0.05, * for 0.01 < p-value ≤ 0.05, ** for 0.001 < p-value ≤ 0.01, *** for 0.0001 < p-value ≤ 0.001).

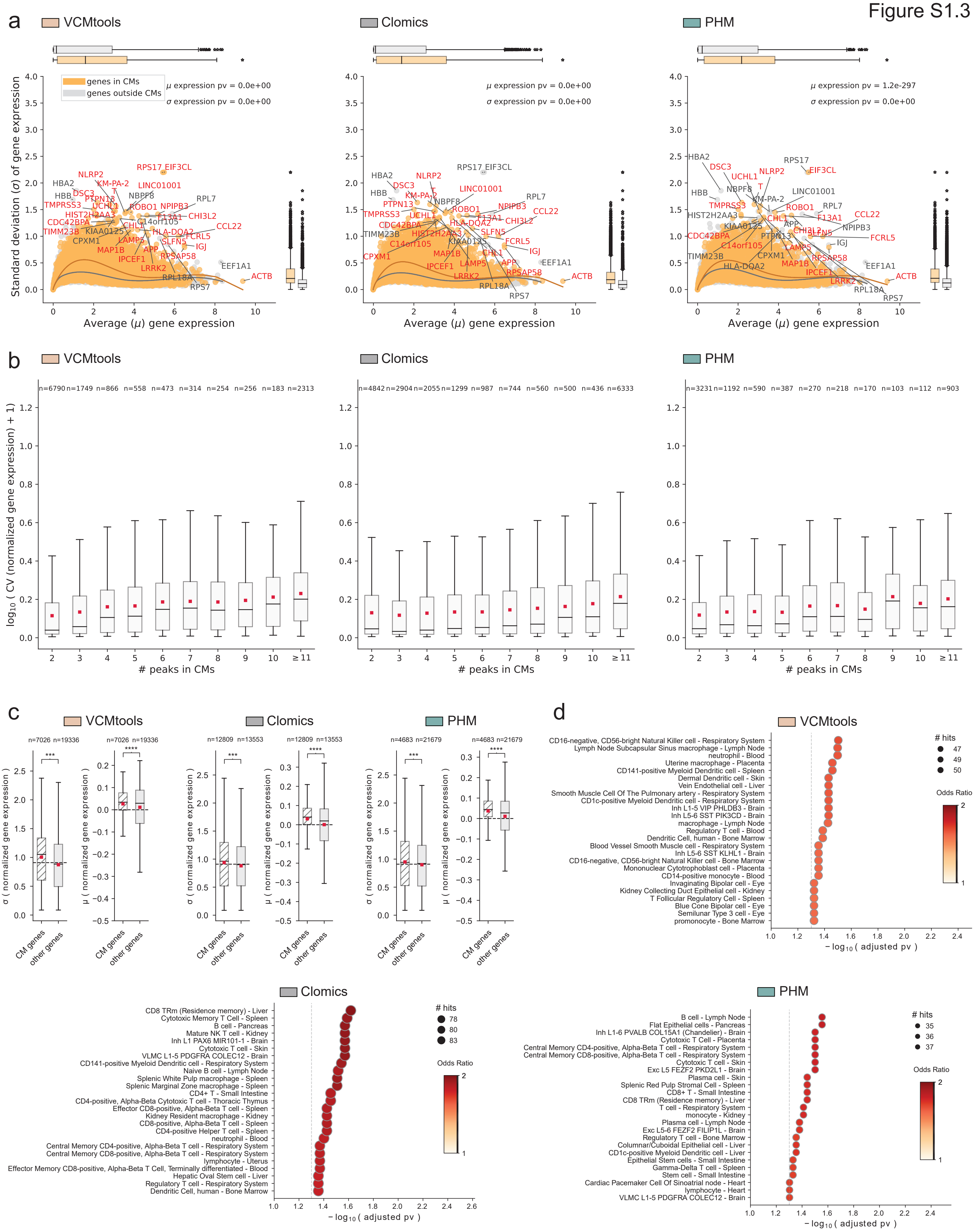
Figure S1.3
a
VCMtools
Clomics
PHM
genes in CMs
genes outside CMs
μ expression pv = 0.0e+00
σ expression pv = 0.0e+00
μ expression pv = 1.2e-297
Standard deviation (σ) of gene expression
Average (μ) gene expression
b
log10 ( CV (normalized gene expression) + 1)
### peaks in CMs
c
σ ( normalized gene expression )
μ ( normalized gene expression )
CM genes
other genes
d
### hits
Odds Ratio
− log10 ( adjusted pv )

**Figure S1.3. CMs mapped with different methods capture relevant genes in a given cell type. a.** ***From left to right***: VCMtools, Clomics, PHM. Average gene expression versus standard deviation of gene expression for all protein-coding genes and lincRNA genes colored by localization: genes overlapped by CMs (in orange), genes outside CMs (in gray). The boxplots at the top of the scatter plot and on the right-hand side show quantiles of the gene expression means and standard deviations respectively by category. P-values (pv) in the upper right corner indicate the Mann-Whitney U p-values for the tests performed between the gene groups for gene expression means ($\mu$) and standard deviations ($\sigma$). **b**. ***From left to right***: VCMtools, Clomics, PHM. Log10 coefficient of variation of normalized gene expression for genes falling into CMs of different sizes. **c.** Standard deviation and mean of normalized gene expression for genes overlapped by CMs (CM genes) and not overlapped by CMs (other genes). **d.** ***From left to right***: VCMtools, Clomics, PHM. Gene Ontology terms for genes overlapped by CMs show enrichment for B-cell-specific annotations. P-value indications are non-significant (ns) for p-value > 0.05, * for 0.01 < p-value ≤ 0.05, ** for 0.001 < p-value ≤ 0.01, *** for 0.0001 < p-value ≤ 0.001, **** p-value ≤ 0.0001.

Figure S2.1

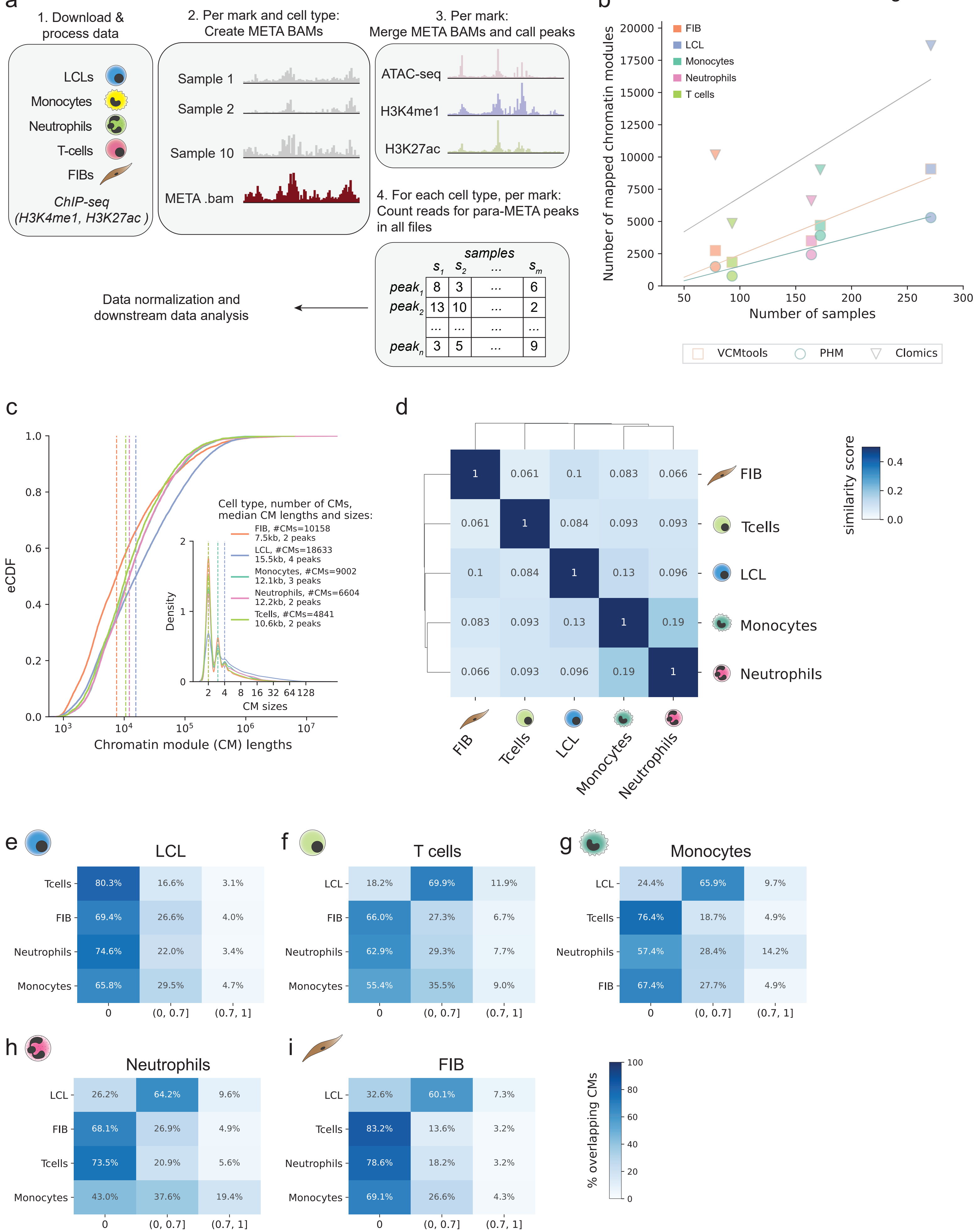

a
1. Download & process data
LCLs
Monocytes
Neutrophils
T-cells
FIBs
ChIP-seq (H3K4me1, H3K27ac )
2. Per mark and cell type: Create META BAMs
Sample 1
Sample 2
Sample 10
META .bam
3. Per mark: Merge META BAMs and call peaks
ATAC-seq
H3K4me1
H3K27ac
4. For each cell type, per mark: Count reads for para-META peaks in all files
samples
s1 s2 ... sm
peak1 8 3 ... 6
peak2 13 10 ... 2
... ... ... ...
peakn 3 5 ... 9
Data normalization and downstream data analysis
b
FIB
LCL
Monocytes
Neutrophils
T cells
Number of mapped chromatin modules
Number of samples
VCMtools
PHM
Clomics
c
eCDF
Chromatin module (CM) lengths
Cell type, number of CMs, median CM lengths and sizes:
FIB, #CMs=10158 7.5kb, 2 peaks
LCL, #CMs=18633 15.5kb, 4 peaks
Monocytes, #CMs=9002 12.1kb, 3 peaks
Neutrophils, #CMs=6604 12.2kb, 2 peaks
Tcells, #CMs=4841 10.6kb, 2 peaks
Density
CM sizes
d
similarity score
FIB
Tcells
LCL
Monocytes
Neutrophils
e
LCL
f
T cells
g
Monocytes
h
Neutrophils
i
FIB
% overlapping CMs
0
(0, 0.7]
(0.7, 1]

**Figure S2.1. CM mapping and comparison across cell types. a.** Schematic representation of the data processing pipeline. First, we downloaded the BAM files for all individuals available for a particular cell type (step 1). The top 10 largest BAM files per ChIP-seq histone modification and per cell type were down-sampled to an equal number of reads and merged to create a "meta"-BAM file by stacking up the signal across the files (step 2). For each histone mark, we merged "meta"-BAM files across cell types and used this final merged meta bam file call peaks (step 3). For each cell type, for each histone mark, we counted reads and created a count matrix from the merged "meta"-BAM file (step 4). Known (based on the available meta data) and unknown covariates (based on principal component analyses) were removed. The resulting count matrices were used for downstream analyses including CM mapping. **b.** Dependency between the number of samples in each cell type and the number of mapped CMs per CM mapping method. Lines show linear fits to the data per method. **c.** Empirical Cumulative Density Function (eCDF) of CM lengths and sizes (inset panels) mapped with Clomics. **d.** Average pairwise similarity scores (harmonic mean (HM) based) between cell types. **e-i.** Percentage of CMs mapped in a cell type (indicated above each heatmap) that overlap with CMs in other cell types at different similarity score ranges: 0 – no overlap, (0-0.7] – partial overlap, (0.7, 1] – high similarity/identical CMs. One-sided comparison of CM similarities in one cell type versus the others shows 43-83.2% of CMs not being captured in other cell types. When comparing CMs mapped in any cell type to CMs identified in LCLs, 60-70% of CMs falling into the partially overlapping CM group, which can be explained by the extensive number of CMs mapped in LCLs as compared to other cell types.

Figure S2.2

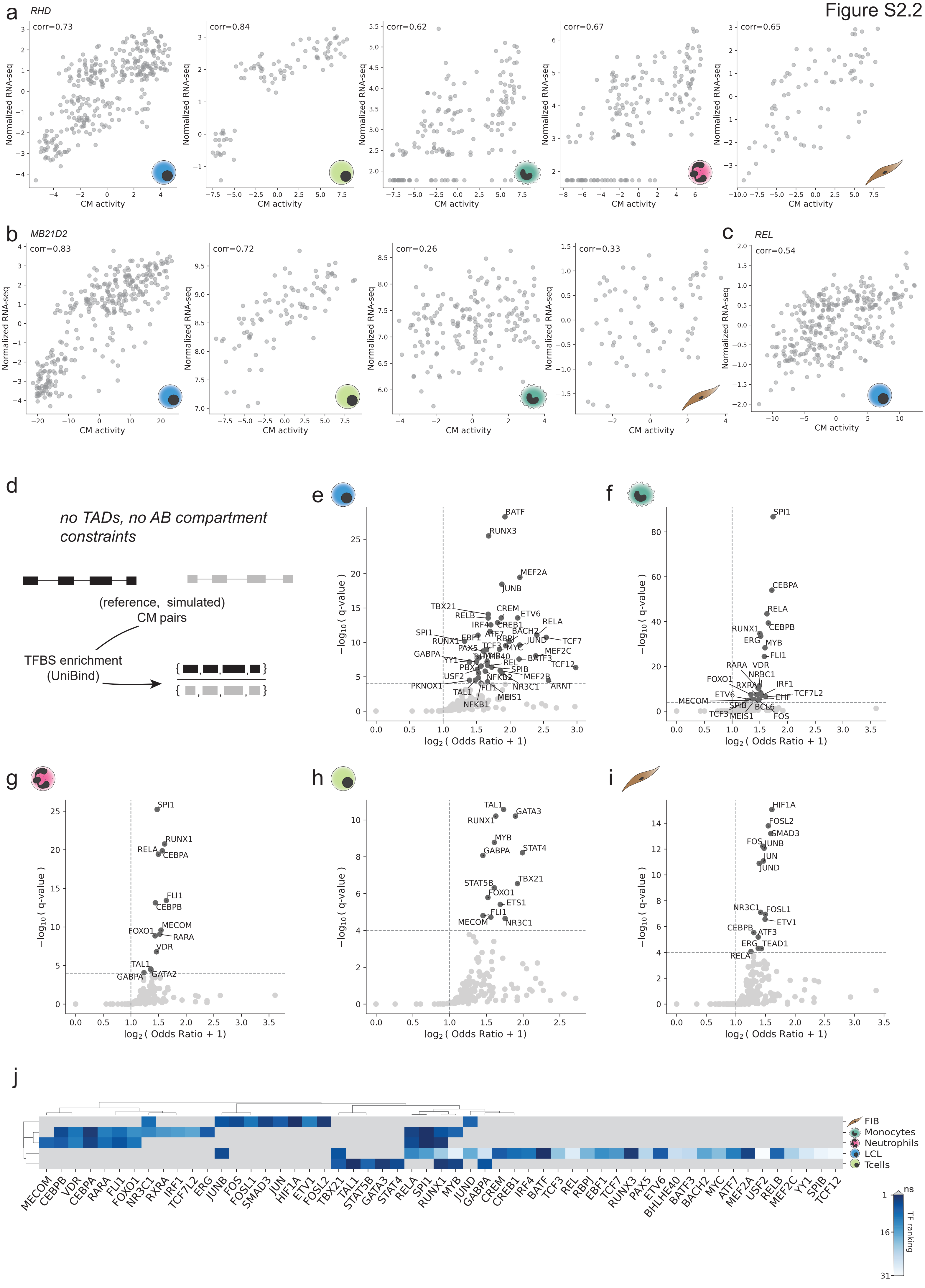
a
RHD
corr=0.73
corr=0.84
corr=0.62
corr=0.67
corr=0.65
Normalized RNA-seq
CM activity
b
MB21D2
corr=0.83
corr=0.72
corr=0.26
corr=0.33
c
REL
corr=0.54
d
no TADs, no AB compartment constraints
(reference, simulated) CM pairs
TFBS enrichment (UniBind)
e
f
g
h
i
−log10 ( q-value )
log2 ( Odds Ratio + 1)
BATF
RUNX3
MEF2A
JUNB
SPI1
CEBPA
RELA
CEBPB
RUNX1
MYB
FLI1
TAL1
GATA3
HIF1A
FOSL2
SMAD3
j
FIB
Monocytes
Neutrophils
LCL
Tcells
TF ranking
ns

**Figure S2.2. CM activity and TF binding associated with CM regions. a-c.** Activity of CMs per cell type (if there is a CM; in the case of several CMs in a locus, the highest correlation between aCM score and RNA-seq data is shown) versus normalized RNA-seq values for *RHD*, *MB21D2* and *REL* genes. **d.** Schematic representation of the differential TFBS enrichment analysis strategy for the paired set of CMs and simulated CMs. Simulated CMs were generated without constraining the genomic reference location to being confined within TADs and A/B compartments. **e-i.** ***From left to right****:* data for LCLs, Monocytes. ***Second row from left to right****:* Neutrophils, T cells, Fibroblasts. TFBS enrichment results per cell type shown as the log2 Odds Ratio versus -log10 q-value of TFBS enrichment within individual cell types when contrasting mapped ("reference") CM peaks vs simulated CM peaks. **j.** Heatmap of TF ranking for TFBSs that passed the q-value threshold (q-value ≤ 1e-04) in at least one of the cell types indicated on the right side. Gray color indicates non-significant hits.

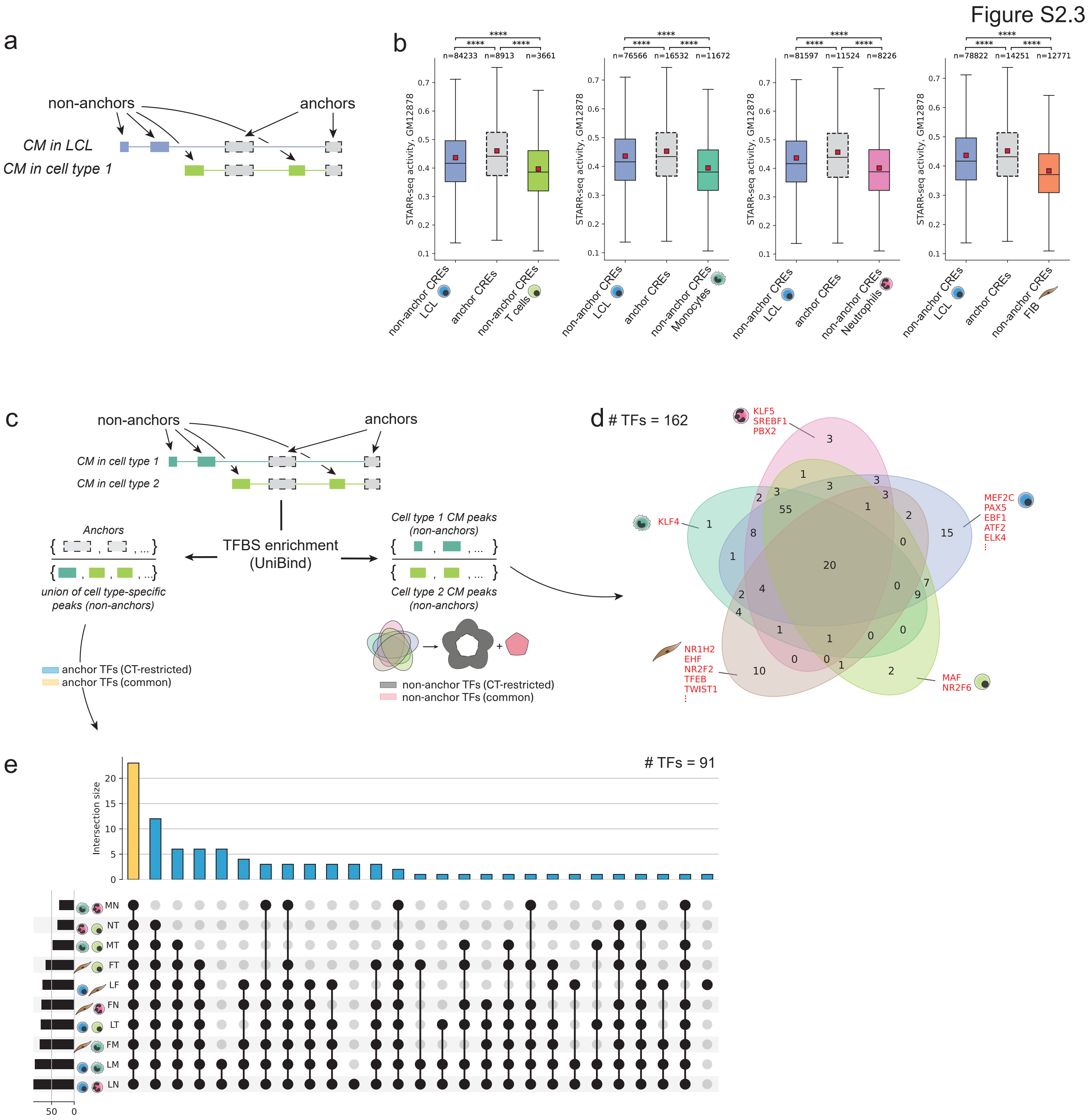
Figure S2.3
a
non-anchors
anchors
CM in LCL
CM in cell type 1
b
STARR-seq activity, GM12878
n=84233 n=8913 n=3661
n=76566 n=16532 n=11672
n=81597 n=11524 n=8226
n=78822 n=14251 n=12771
non-anchor CREs LCL
anchor CREs
non-anchor CREs T cells
non-anchor CREs Monocytes
non-anchor CREs Neutrophils
non-anchor CREs FIB
c
non-anchors
anchors
CM in cell type 1
CM in cell type 2
Anchors
union of cell type-specific peaks (non-anchors)
TFBS enrichment (UniBind)
Cell type 1 CM peaks (non-anchors)
Cell type 2 CM peaks (non-anchors)
anchor TFs (CT-restricted)
anchor TFs (common)
non-anchor TFs (CT-restricted)
non-anchor TFs (common)
d
### TFs = 162
KLF5
SREBF1
PBX2
KLF4
MEF2C
PAX5
EBF1
ATF2
ELK4
NR1H2
EHF
NR2F2
TFEB
TWIST1
MAF
NR2F6
e
### TFs = 91
Intersection size
MN
NT
MT
FT
LF
FN
LT
FM
LM
LN

**Figure S2.3. TFBS- and STARR-seq-based classification based of anchor/non-anchor CM elements. a.** Schematic representation of how CM CREs are categorized into groups of anchors (shared) and non-anchor (cell type-restricted) CREs in one comparison between two cell types. **b**. Activity of CM CREs, as measured with STARR-seq in GM12878, for pairwise comparisons of LCLs and all other cell types (***from left to right***: T cells, Monocytes, Neutrophils, FIB). Individual side-wise boxplots per panel correspond to STARR-seq activities at cell type-specific CREs for a given cell type pair, whereas the central boxplot shows activity at anchor CM CREs that are thus shared between the respective cell types. **c.** Schematic representation of the differential TFBS enrichment analysis strategy for CM peaks across all pairwise cell type comparisons. First, for a given cell type pair, we split CM CREs into sets of anchor and non-anchor CREs. Next, we performed differential TFBS enrichment analysis with UniBind for 1) anchor vs non-anchor (union of cell type-specific) CREs, 2) non-anchor CREs in cell type 1 vs non-anchor CREs in cell type 2. Then, we split anchor and non-anchor CREs into cell type (CT)-restricted (TFBSs enriched in one or a few cell types) and CT-common groups (TFBSs enriched in all cell types). **d.** Venn diagram revealing the overlaps between enrichments within non-anchor CREs in a cell type-specific manner. TFBS enrichment in non-anchor CM CREs of one cell type vs non-anchor CM CREs in all other cell types. Significantly enriched TFBSs (q-value ≤ 1e-06) obtained through this comparison were additionally filtered based on TF expression (pTPM > 2 in at least one cell type), resulting in a total of 162 TF candidates. **e.** UpSet plot for TFBS enrichment results when contrasting anchor and non-anchor CREs. The total number of TFs that are considered here is 91.

Figure S2.4

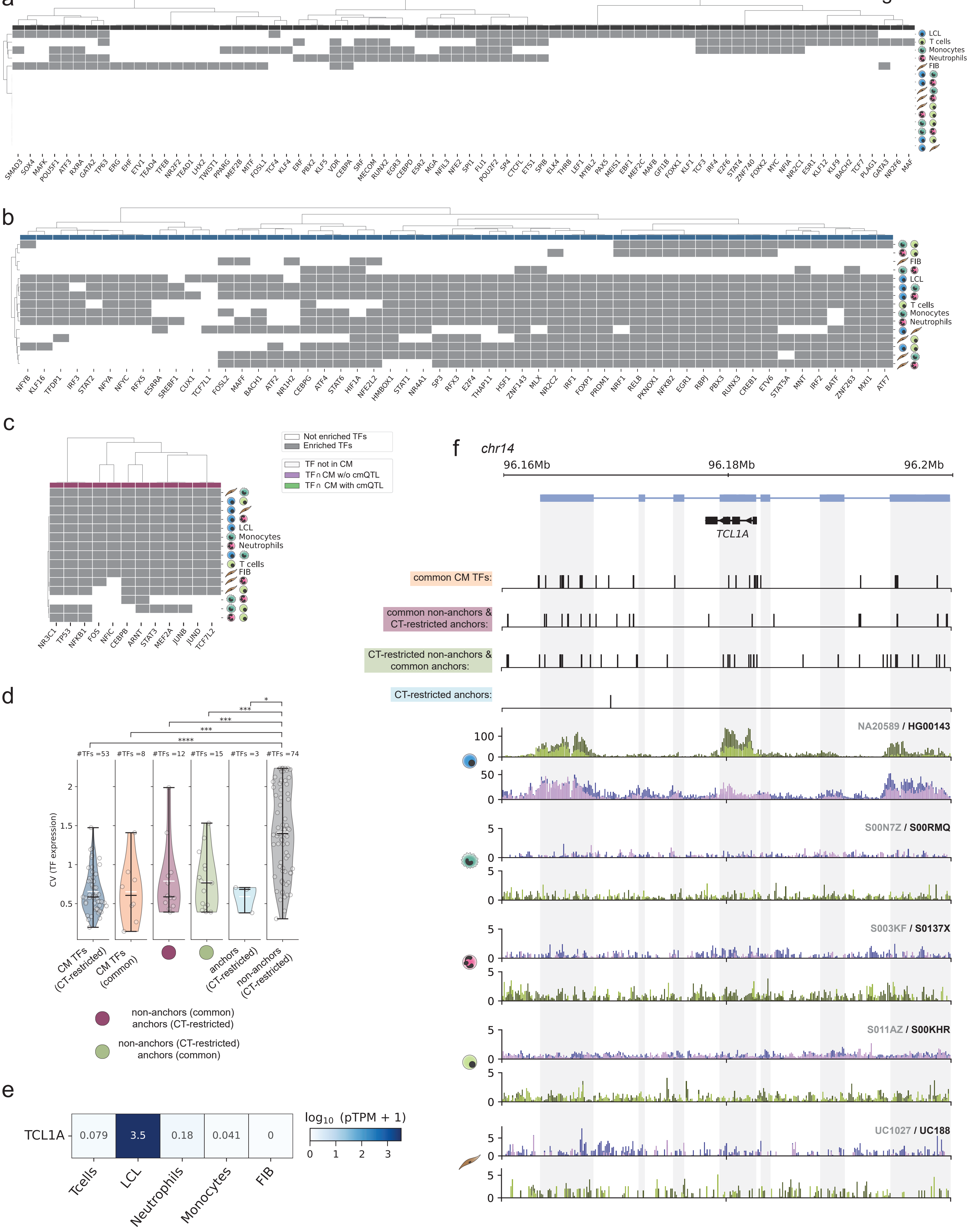

a
LCL
T cells
Monocytes
Neutrophils
FIB
SMAD3 SOX4 MAFK POU5F1 ATF3 RXRA GATA2 TP63 ERG EHF ETV1 TEAD4 TFEB NR2F2 TEAD1 LHX2 TWIST1 PPARG MEF2B MITF FOSL1 TCF4 KLF4 ERF PBX2 KLF5 VDR CEBPA SRF MECOM RUNX2 EGR3 CEBPD ESR2 MGA NFIL3 NFE2 SPI1 FLI1 POU2F2 SP4 CTCFL ETS1 SPIB ELK4 THRB LEF1 MYBL2 PAX5 MEIS1 EBF1 MEF2C MAFB GFI1B FOXK1 KLF1 TCF3 IRF4 E2F6 STAT4 ZNF740 FOXK2 MYC NFIA NR2C1 ESR1 KLF12 KLF9 BACH2 TCF7 PLAG1 GATA3 NR2F6 MAF
b
FIB
LCL
T cells
Monocytes
Neutrophils
NFYB KLF16 TFDP1 IRF3 STAT2 NFYA NFYC RFX5 ESRRA SREBF1 CUX1 TCF7L1 FOSL2 MAFF BACH1 ATF2 NR1H2 CEBPG ATF4 STAT6 HIF1A NFE2L2 HMBOX1 STAT1 NR4A1 SP3 RFX3 E2F4 THAP11 HSF1 ZNF143 MLX NR2C2 IRF1 FOXP1 PRDM1 NRF1 RELB PKNOX1 NFKB2 EGR1 RBPJ PBX3 RUNX3 CREB1 ETV6 STAT5A MNT IRF2 BATF ZNF263 MXI1 ATF7
c
Not enriched TFs
Enriched TFs
TF not in CM
TF∩CM w/o cmQTL
TF∩ CM with cmQTL
LCL
Monocytes
Neutrophils
T cells
FIB
NR3C1 TP53 NFKB1 FOS NFIC CEBPB ARNT STAT3 MEF2A JUNB JUND TCF7L2
d
#TFs =53
#TFs =8
#TFs =12
#TFs =15
#TFs =3
#TFs =74
****
***
***
***
*
CV (TF expression)
CM TFs (CT-restricted)
CM TFs (common)
anchors (CT-restricted)
non-anchors (CT-restricted)
non-anchors (common)
anchors (CT-restricted)
non-anchors (CT-restricted)
anchors (common)
e
TCL1A
0.079
3.5
0.18
0.041
0
Tcells
LCL
Neutrophils
Monocytes
FIB
log10 (pTPM + 1)
0
1
2
3
f
chr14
96.16Mb
96.18Mb
96.2Mb
TCL1A
common CM TFs:
common non-anchors &
CT-restricted anchors:
CT-restricted non-anchors &
common anchors:
CT-restricted anchors:
NA20589 / HG00143
S00N7Z / S00RMQ
S003KF / S0137X
S011AZ / S00KHR
UC1027 / UC188

**Figure S2.4. Binding sites from different TF classes show differential enrichment in a cell type-dependent manner. a-c. *From top to bottom***: cell type (CT)-restricted non-anchors (dark gray; **a**), CT-restricted CM TFs (dark blue, **b**), CT-restricted anchors and common non-anchors (maroon, **c**). Boolean heatmap of TFBS enriched (gray for enriched TFBS, white for non-enriched) in the group per cell type or cell type pair. **d.** Coefficient of variation of TF expression (pTPM) between cell types for subclasses of TFs identified through differential TFBS enrichment analysis. Stars indicate the respective p-value strength for the Mann-Whitney U test. **e.** *TCL1A* expression (pTPM) across cell types. **f. *Left***: Tracks with black vertical lines indicate TFBSs for TFs falling into defined categories (see **Fig 2** for the respective legend). The bottom tracks correspond to ChIP-seq profiles of H3K27ac and H3K4me1 for two individuals, one with the highest (dark blue (H3K4me1) and dark green (H3K27ac)) and one with the lowest (magenta (H3K4me1) and lime color (H3K27ac)) CM activity in the *TCL1A* gene locus for all cell types. ***From top to bottom***: LCLs, Monocytes, Neutrophils, T cells, FIB. P-value indications are non-significant (ns) for p-value > 0.05, * for 0.01 < p-value ≤ 0.05, ** for 0.001 < p-value ≤ 0.01, *** for 0.0001 < p-value ≤ 0.001, **** p-value ≤ 0.0001.

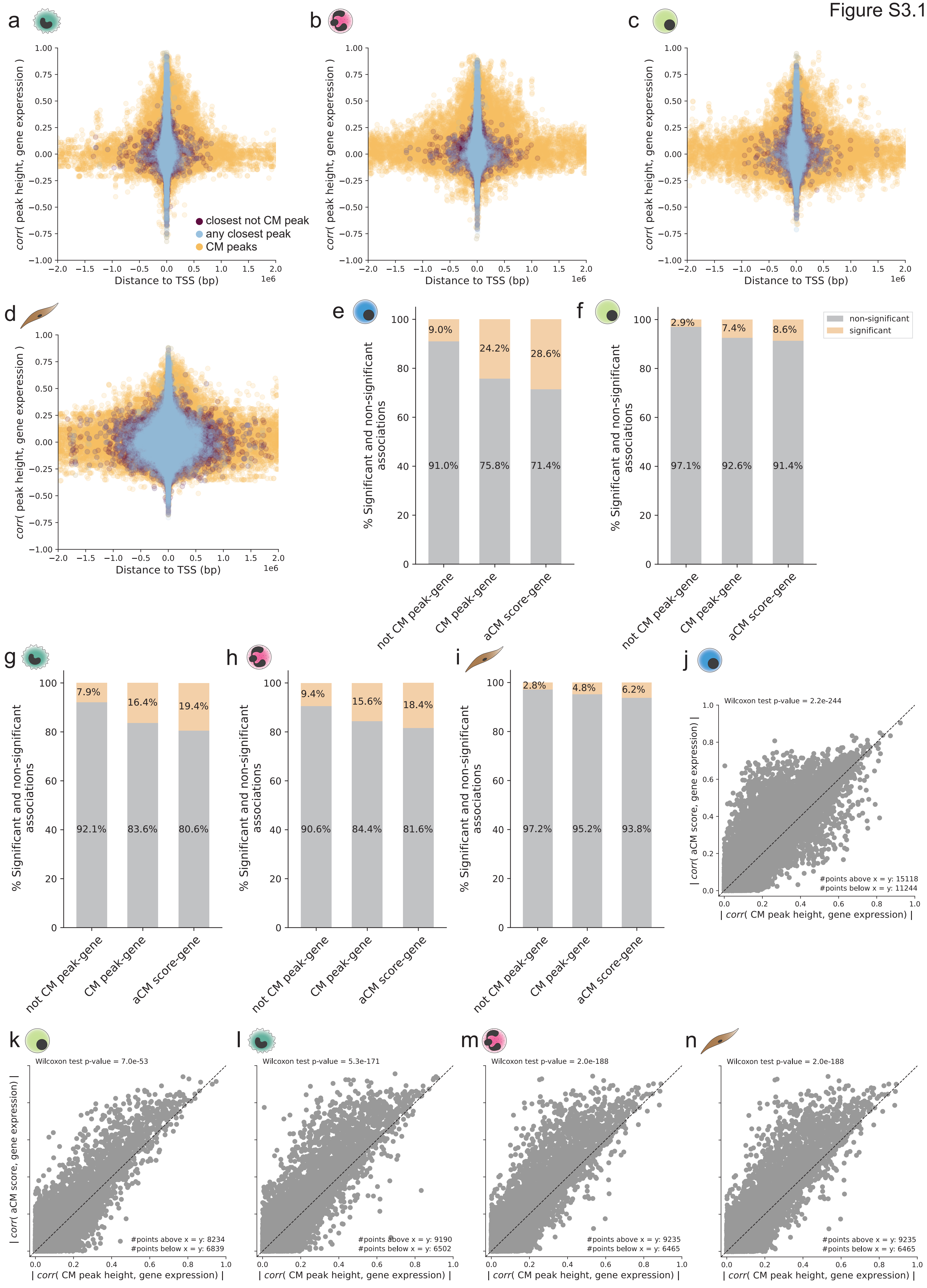
a
b
c
d
e
f
g
h
i
j
k
l
m
n
corr( peak height, gene experession )
Distance to TSS (bp)
1e6
closest not CM peak
any closest peak
CM peaks
% Significant and non-significant associations
non-significant
significant
not CM peak-gene
CM peak-gene
aCM score-gene
9.0%
24.2%
28.6%
91.0%
75.8%
71.4%
2.9%
7.4%
8.6%
97.1%
92.6%
91.4%
7.9%
16.4%
19.4%
92.1%
83.6%
80.6%
9.4%
15.6%
18.4%
90.6%
84.4%
81.6%
2.8%
4.8%
6.2%
97.2%
95.2%
93.8%
Wilcoxon test p-value = 2.2e-244
#points above x = y: 15118
#points below x = y: 11244
Wilcoxon test p-value = 7.0e-53
#points above x = y: 8234
#points below x = y: 6839
Wilcoxon test p-value = 5.3e-171
#points above x = y: 9190
#points below x = y: 6502
Wilcoxon test p-value = 2.0e-188
#points above x = y: 9235
#points below x = y: 6465
Wilcoxon test p-value = 2.0e-188
#points above x = y: 9235
#points below x = y: 6465
| corr( aCM score, gene expression) |
| corr( CM peak height, gene expression) |

**Figure S3.1. CMs and CM activity explain gene regulatory variation better than individual peaks. a-d.** ***From left to right***: Monocytes, Neutrophils, T cells, Fibroblasts. Distance to the closest gene transcription start site (TSS) from the peak center versus correlation between the peak height and the closest gene expression. The colors indicate different peak categories, where "any peak" group corresponds to the closest peak to the TSS irrespective of its annotation as CM peak or non-CM peak. **e-i.** ***From left to right***: LCLs, T cells, Monocytes, Neutrophils, FIB. Number of significant (light orange) and non-significant (gray) peak-gene associations based on the correlation for three tested groups ***from left to right***: non-CM peak to gene, CM peak to gene, and aCM score to gene**. j-n.** ***From left to right***: LCLs, T cells, Monocytes, Neutrophils, FIB. Absolute correlation values between height of a peak belonging to a CM and closest gene expression versus absolute correlation values between aCM score and closest gene expression. Wilcoxon test p-value for the differences between the values on the x-axis and the values on the y-axis are indicated in the upper left corner.

Figure S3.2

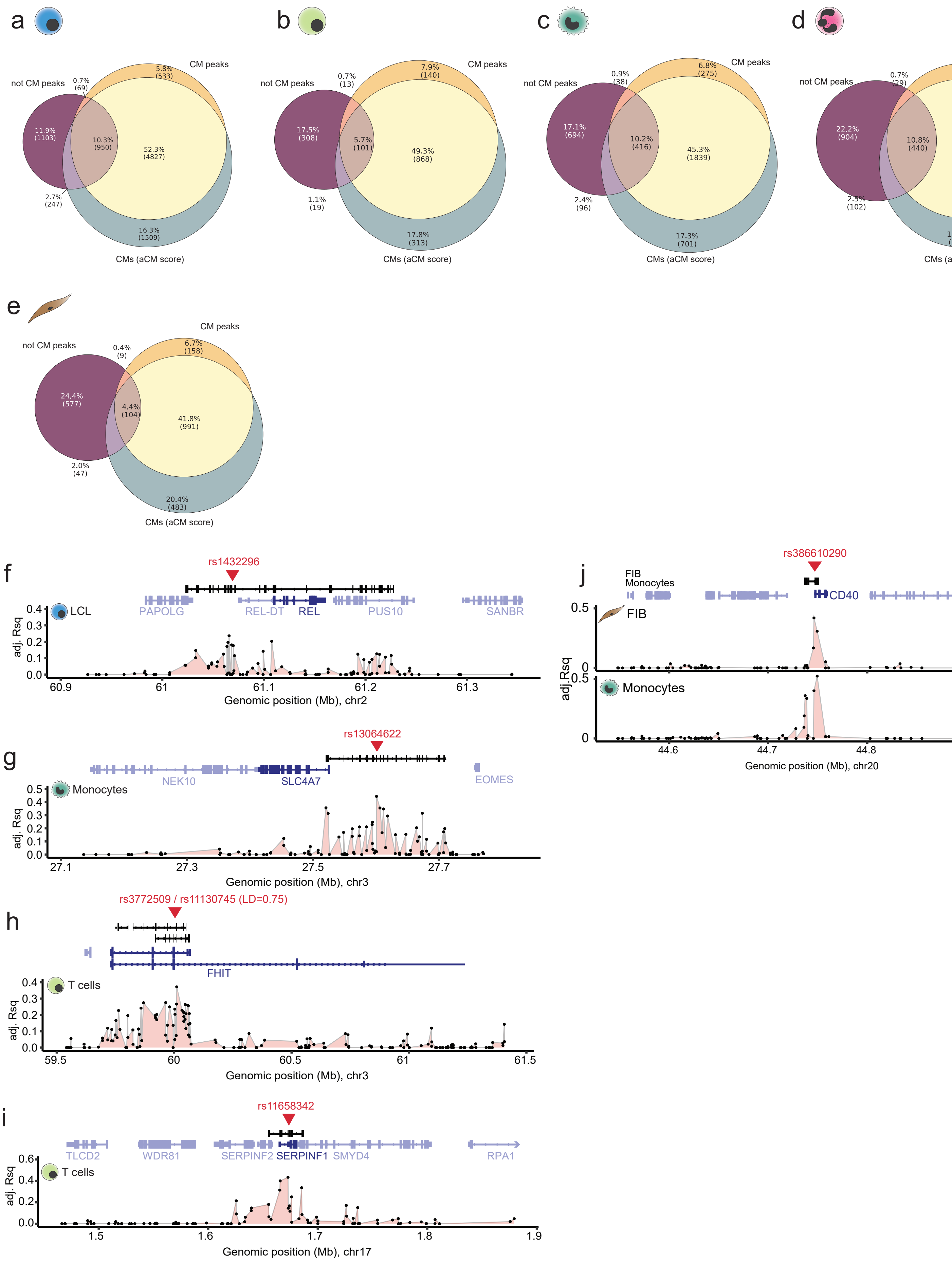

a
not CM peaks
CM peaks
CMs (aCM score)
b
c
d
e
f
rs1432296
LCL
PAPOLG
REL-DT
REL
PUS10
SANBR
adj. Rsq
Genomic position (Mb), chr2
g
rs13064622
Monocytes
NEK10
SLC4A7
EOMES
Genomic position (Mb), chr3
h
rs3772509 / rs11130745 (LD=0.75)
T cells
FHIT
Genomic position (Mb), chr3
i
rs11658342
T cells
TLCD2
WDR81
SERPINF2
SERPINF1
SMYD4
RPA1
Genomic position (Mb), chr17
j
rs386610290
FIB
Monocytes
CD40
Genomic position (Mb), chr20

**Figure S3.2. cmQTLs often impact the strongest CM peak. a-e.** ***From left to right***: LCL, T cells, Monocytes, Neutrophils, FIB. Venn diagram indicating the percentages of genes falling into different significant peak-gene or aCM-gene association categories. **f-i.** Examples of CMs spanning genes in various cell types having a cmQTL (red triangle). The tracks below CMs and genes show the association strength (adjusted $R^2$ of the linear regression) between every peak in the locus and expression of the gene highlighted in dark blue.

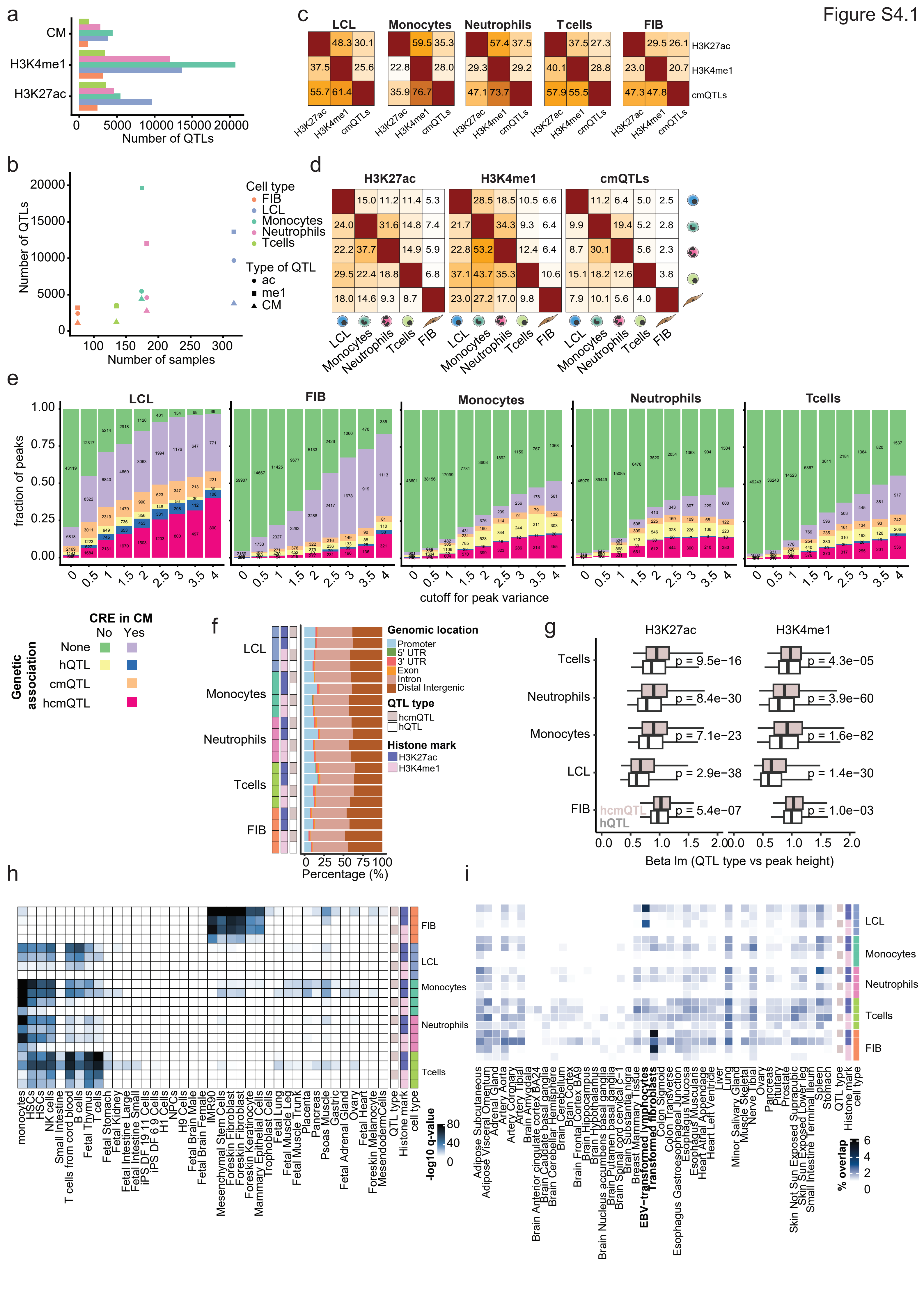
a
CM
H3K4me1
H3K27ac
Number of QTLs
c
LCL
Monocytes
Neutrophils
T cells
FIB
H3K27ac
H3K4me1
cmQTLs
b
Number of QTLs
Number of samples
Cell type
FIB
LCL
Monocytes
Neutrophils
Tcells
Type of QTL
ac
me1
CM
d
H3K27ac
H3K4me1
cmQTLs
e
fraction of peaks
cutoff for peak variance
CRE in CM
No
Yes
Genetic association
None
hQTL
cmQTL
hcmQTL
f
Genomic location
Promoter
5' UTR
3' UTR
Exon
Intron
Distal Intergenic
QTL type
hcmQTL
hQTL
Histone mark
H3K27ac
H3K4me1
Percentage (%)
g
p = 9.5e−16
p = 4.3e−05
p = 8.4e−30
p = 3.9e−60
p = 7.1e−23
p = 1.6e−82
p = 2.9e−38
p = 1.4e−30
p = 5.4e−07
p = 1.0e−03
Beta lm (QTL type vs peak height)
h
-log10 q-value
i
% overlap
QTL type
Histone mark
cell type

**Figure S4.1. Categorization of cmQTLs and hQTLs. a.** Total number of QTLs mapped per category. **b.** Relation between the number of mapped QTLs and number of included individuals. **c.** Percentage of QTLs that are also in LD ($R^2 > 0.8$) with another type of QTL. **d.** Percentages of QTLs shared (or in LD ($R^2 > 0.8$)) between cell types. **e.** H3K27ac peaks were binned by degree of interindividual variance. Numbers on the x-axis represent the number of standard deviations (SD) by which the peaks are variable. For example, 0 is the bin 0-0.5 SD, 0.5 indicates the bin 0.5-1 SD, and 4 represents more than 4 SD variable. Each number on the plot represents the number of peaks in each category. The y-axis represents the cumulative percentage per bin. **f.** Histone QTLs were divided in two groups: histone QTL only (hQTL) and histone + cmQTL (hcmQTL). Genomic locations of the variants in each group are shown. **g.** Boxplots showing the beta values versus height of the peak that the histone QTL affects. P-values were calculated using a Wilcoxon test. **h.** Enrichment of variants in open chromatin regions of each of the indicated cell types. A higher -log10 p-value indicates stronger enrichment. **i.** Percentage of variants overlapping or in LD ($R^2 > 0.8$) with eQTLs in different GTEx tissues.

Figure S4.2

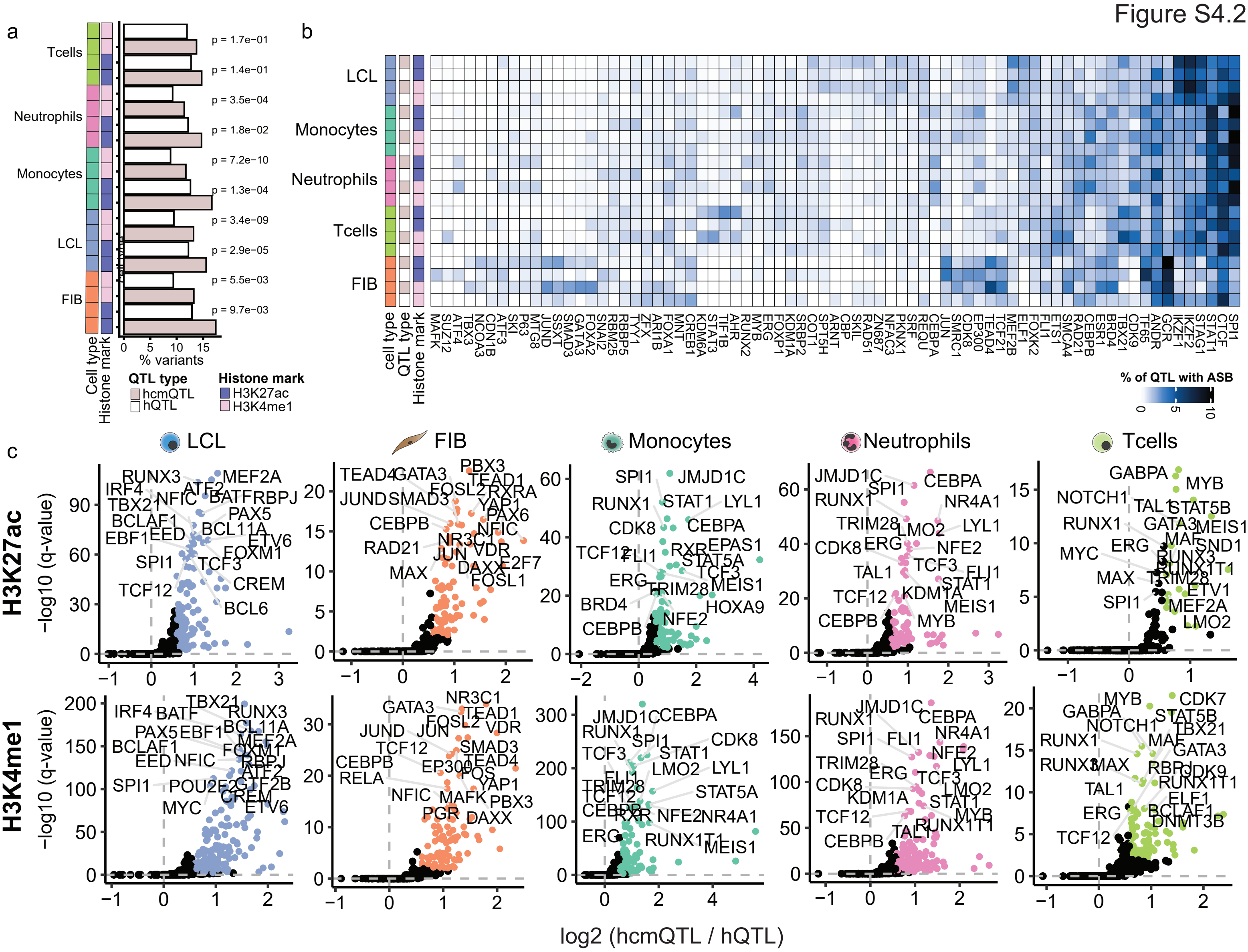
a
Tcells
Neutrophils
Monocytes
LCL
FIB
p = 1.7e−01
p = 1.4e−01
p = 3.5e−04
p = 1.8e−02
p = 7.2e−10
p = 1.3e−04
p = 3.4e−09
p = 2.9e−05
p = 5.5e−03
p = 9.7e−03
% variants
Cell type
Histone mark
QTL type
hcmQTL
hQTL
Histone mark
H3K27ac
H3K4me1
b
LCL
Monocytes
Neutrophils
Tcells
FIB
cell type
QTL type
Histone mark
% of QTL with ASB
c
LCL
FIB
Monocytes
Neutrophils
Tcells
H3K27ac
H3K4me1
−log10 (q-value)
log2 (hcmQTL / hQTL)

**Figure S4.2. hcmQTLs enrich for binding of cell type-specific TFs. a.** Percentage of genetic variants with at least one allele-specific binding (ASB) event. P-values were calculated using a Fisher's exact test. **b.** Heatmap indicating the percentage of variants that have an ASB associated with each of the indicated TFs. **c.** Scatter plots showing the log2 enrichment of TF binding in 200 bp windows around hcmQTLs compared to hQTLs. Q-values were obtained using a Benjamini-Hochberg correction of p-values associated with the enrichment.

Figure S4.3

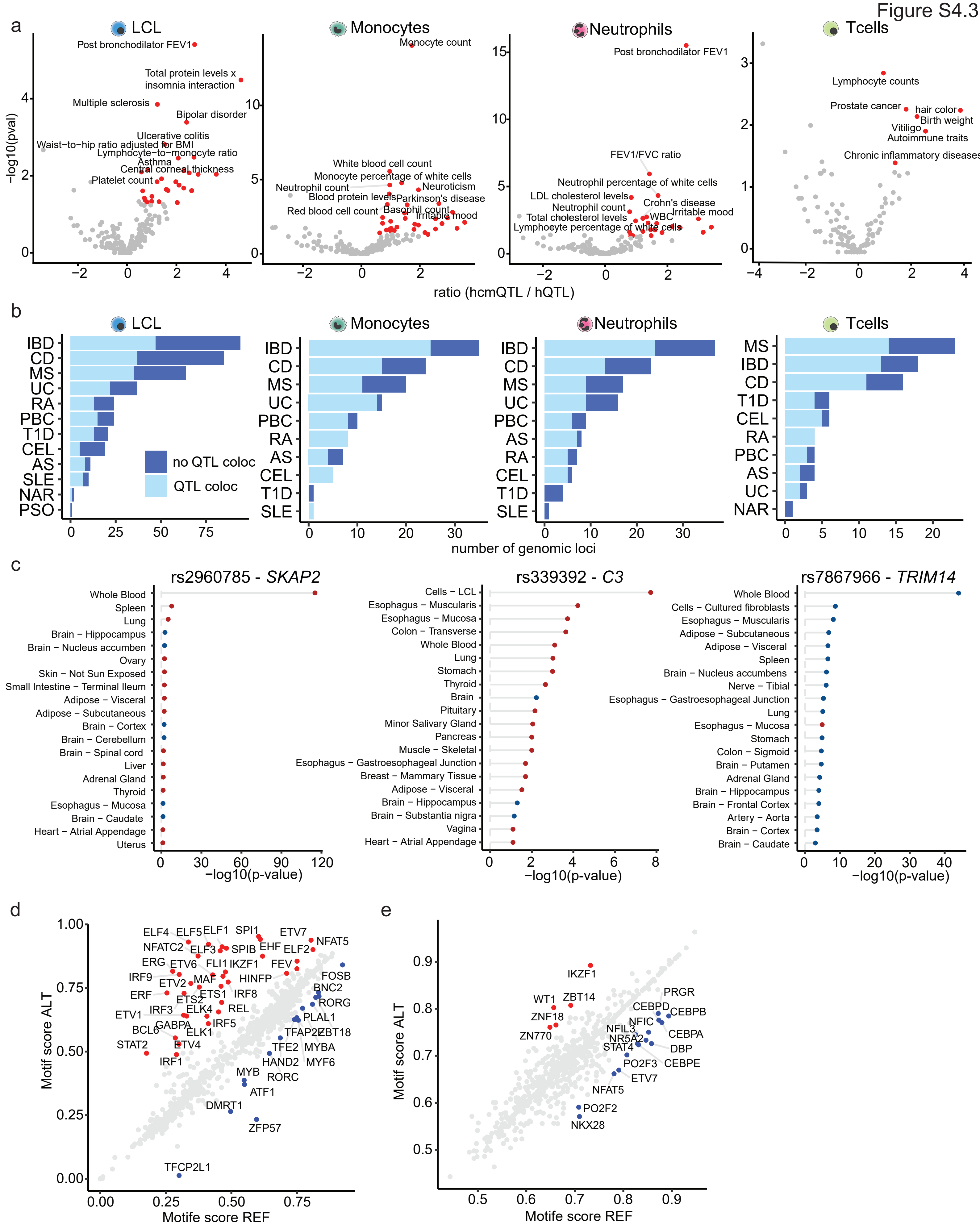

a
LCL
Monocytes
Neutrophils
Tcells
-log10(pval)
ratio (hcmQTL / hQTL)
Post bronchodilator FEV1
Total protein levels x insomnia interaction
Multiple sclerosis
Bipolar disorder
Ulcerative colitis
Waist−to−hip ratio adjusted for BMI
Lymphocyte−to−monocyte ratio
Asthma
Central corneal thickness
Platelet count
Monocyte count
White blood cell count
Monocyte percentage of white cells
Neutrophil count
Neuroticism
Blood protein levels
Parkinson's disease
Red blood cell count
Basophil count
Irritable mood
FEV1/FVC ratio
Neutrophil percentage of white cells
LDL cholesterol levels
Crohn's disease
Neutrophil count
Total cholesterol levels
WBC
Lymphocyte percentage of white cells
Lymphocyte counts
Prostate cancer
hair color
Birth weight
Vitiligo
Autoimmune traits
Chronic inflammatory diseases
b
no QTL coloc
QTL coloc
IBD
CD
MS
UC
RA
PBC
T1D
CEL
AS
SLE
NAR
PSO
number of genomic loci
c
rs2960785 - SKAP2
rs339392 - C3
rs7867966 - TRIM14
Whole Blood
Spleen
Lung
Brain − Hippocampus
Brain − Nucleus accumben
Ovary
Skin − Not Sun Exposed
Small Intestine − Terminal Ileum
Adipose − Visceral
Adipose − Subcutaneous
Brain − Cortex
Brain − Cerebellum
Brain − Spinal cord
Liver
Adrenal Gland
Thyroid
Esophagus − Mucosa
Brain − Caudate
Heart − Atrial Appendage
Uterus
Cells − LCL
Esophagus − Muscularis
Colon − Transverse
Stomach
Brain
Pituitary
Minor Salivary Gland
Pancreas
Muscle − Skeletal
Esophagus − Gastroesophageal Junction
Breast − Mammary Tissue
Brain − Substantia nigra
Vagina
Cells − Cultured fibroblasts
Brain − Nucleus accumbens
Nerve − Tibial
Colon − Sigmoid
Brain − Putamen
Brain − Frontal Cortex
Artery − Aorta
−log10(p-value)
d
e
Motif score ALT
Motife score REF
ELF4 ELF5 ELF1 SPI1 ETV7 NFAT5 NFATC2 ELF3 SPIB EHF ELF2 ERG ETV6 FLI1 IKZF1 FEV FOSB IRF9 MAF HINFP BNC2 ETV2 ERF ETS1 IRF8 RORG ETS2 IRF3 ELK4 REL ETV1 PLAL1 GABPA IRF5 BCL6 TFAP2 ZBT18 ELK1 STAT2 ETV4 TFE2 MYBA HAND2 MYF6 IRF1 MYB RORC ATF1 DMRT1 ZFP57 TFCP2L1
IKZF1 PRGR WT1 ZBT14 CEBPD CEBPB ZNF18 NFIC ZN770 NFIL3 CEBPA NR5A2 STAT4 DBP PO2F3 CEBPE ETV7 NFAT5 PO2F2 NKX28

**Figure S4.3. Mapping cell type-specific disruption of epigenome organization by GWAS QTLs using CMs. a.** Log2 observed / expected ratio of hcmQTLs compared to hQTLs for overlap or in LD ($R2 > 0.8$) with GWAS variants. P-values were calculated using a Fisher's exact test. All significant observations ($p < 0.05$) are indicated in red. **b.** Number of autoimmune risk loci with a CM. Color represents whether the candidate cmQTL colocalizes with the GWAS signal at these loci (posterior probability > 0.8 and at least 1 variant that has a p-value of 1e-5 for both GWAS and variant-aCM association). Abbreviations are: spondylitis (AS), Celiac Disease (CEL), Crohn's Disease (CD), Juvenile dermatomyositis (DM), Inflammatory Bowel Disease (IBD), Multiple Sclerosis (MS), primary biliary cirrhosis (PBC), psoriasis (PSO), Rheumatoid Arthritis (RA), Systemic Lupus Erythematosus (SLE), Type 1 Diabetes (T1D) and Ulcerative Colitis (UC). **c.** P-values of associations of rs2960785, rs339392 and rs7867966 with gene expression of *SKAP2*, *C3* and *TRIM14*, respectively, in different sample types derived from the GTEx catalogue. Colours represent whether the minor allele results in increased (red) or decreased (blue) gene expression. **d-e.** Predicted impact of rs339392 (**d**) and rs7867966 (**e**) on the TF motif score (scaled 0 (no match) to 1 (perfect match)) [102].

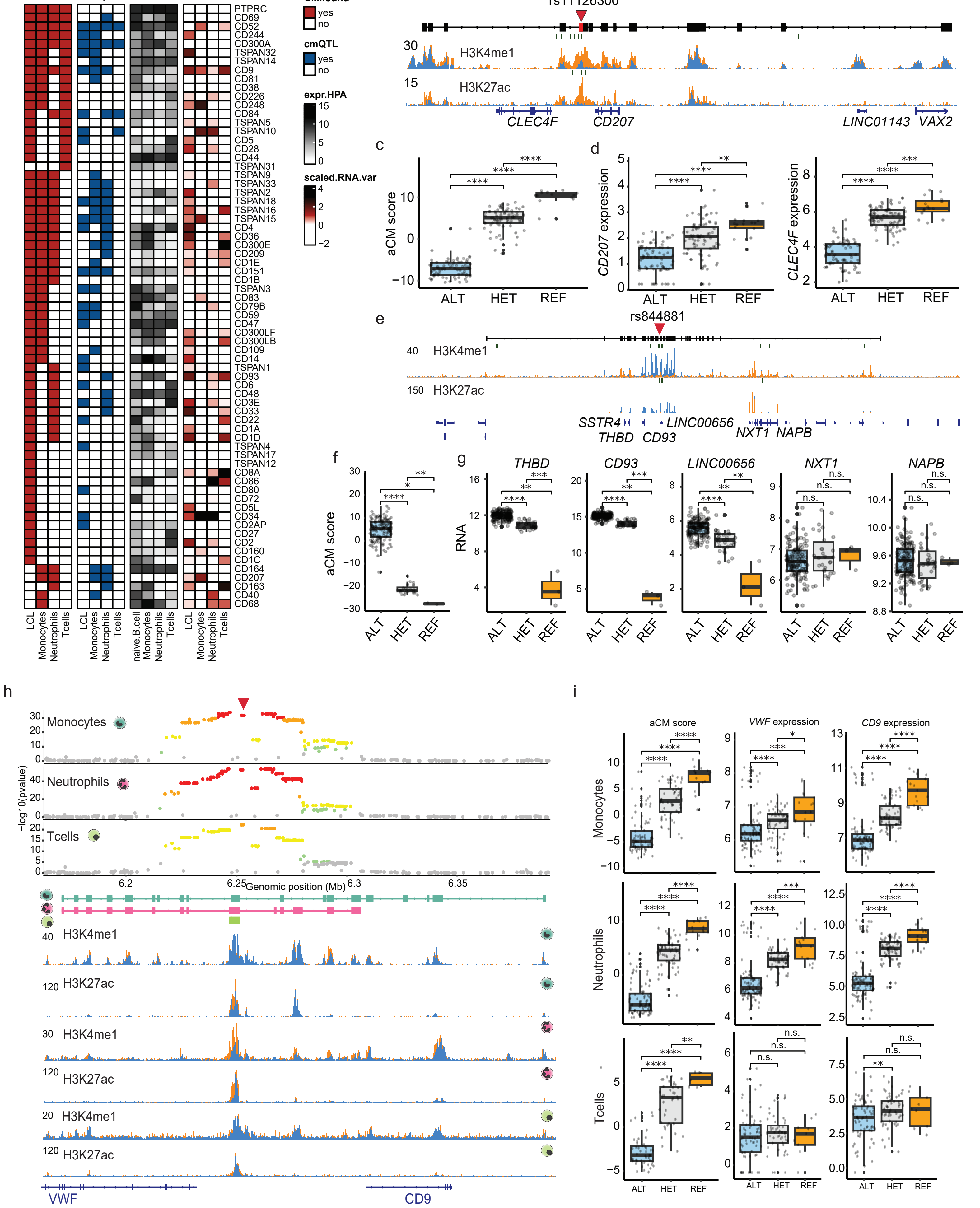
a
CM
cmQTL
HPA
RNA var
CM.found
yes
no
cmQTL
yes
no
expr.HPA
scaled.RNA.var
LCL
Monocytes
Neutrophils
Tcells
naive.B.cell
b
rs11126300
H3K4me1
H3K27ac
CLEC4F
CD207
LINC01143
VAX2
c
aCM score
ALT
HET
REF
d
CD207 expression
CLEC4F expression
e
rs844881
SSTR4
THBD
CD93
LINC00656
NXT1
NAPB
f
g
RNA
n.s.
h
Monocytes
Neutrophils
Tcells
−log10(pvalue)
Genomic position (Mb)
VWF
CD9
i
aCM score
VWF expression
CD9 expression

**Figure S4.4. Genetic variants impacting the epigenome layout around surface receptors through CMs. a.** Heatmap showing surface markers for which a CM was mapped on the genomic locus (column CM), whether a cmQTL was associated (column cmQTL), the normalized protein levels of these genes in different immune cells based on the Human Protein Atlas (column HPA) and the degree of interindividual variation in gene expression (column RNA var). **b.** Example depicting the *CD207* locus in Monocytes, one example individual per reference (orange) and alternative (blue) genotype. The red triangle indicates the cmQTL and green stripes indicate locations where hQTLs were mapped. **c.** Boxplot of the aCM score stratified on the rs11126300 genotype. **d.** Boxplot of *CD207* and *CLEC4F* expression stratified on the rs11126300 genotype. **e.** Example depicting the *CD93* locus in Monocytes, with one example individual per reference (orange) and alternative (blue) genotype. The red triangle indicates the cmQTL and green stripes indicate locations where hQTLs were mapped. **f.** Boxplot of the aCM score stratified on the rs884881 genotype. **g.** Boxplot of gene expression stratified on the rs884881 genotype. Note that expression of *THBD*, *CD93* and *LINC00656*, which are embedded in the CM, follow the aCM pattern, while the genes *NXT1* and *NAPB* that are not embedded in the CM are similarly expressed between rs884881 genotypes. **h.** Example depicting the *VWF* – *CD9* locus in Monocytes, Neutrophils and T cells. The mapped CM in T cells is small, whereas the CM is extended and covers also putative CREs in *VWF* and *CD9* in Monocytes and Neutrophils. While the same variants seem associated with the shared CM region, there is only an impact on gene expression in Monocytes and Neutrophils. One representative individual for the ALT genotype (blue) and the REF genotype (orange) is shown. Note that all ALT and REF tracks originate from cells from the same individual. **i.** Boxplot of the aCM score and gene expression stratified on the genotype of the shared top variant (indicated by the red triangle in (**h**)). P-value indications are non-significant (ns) for p-value > 0.05, * for 0.01 < p-value ≤ 0.05, ** for 0.001 < p-value ≤ 0.01, *** for 0.0001 < p-value ≤ 0.001, **** p-value ≤ 0.0001.

Figure S5

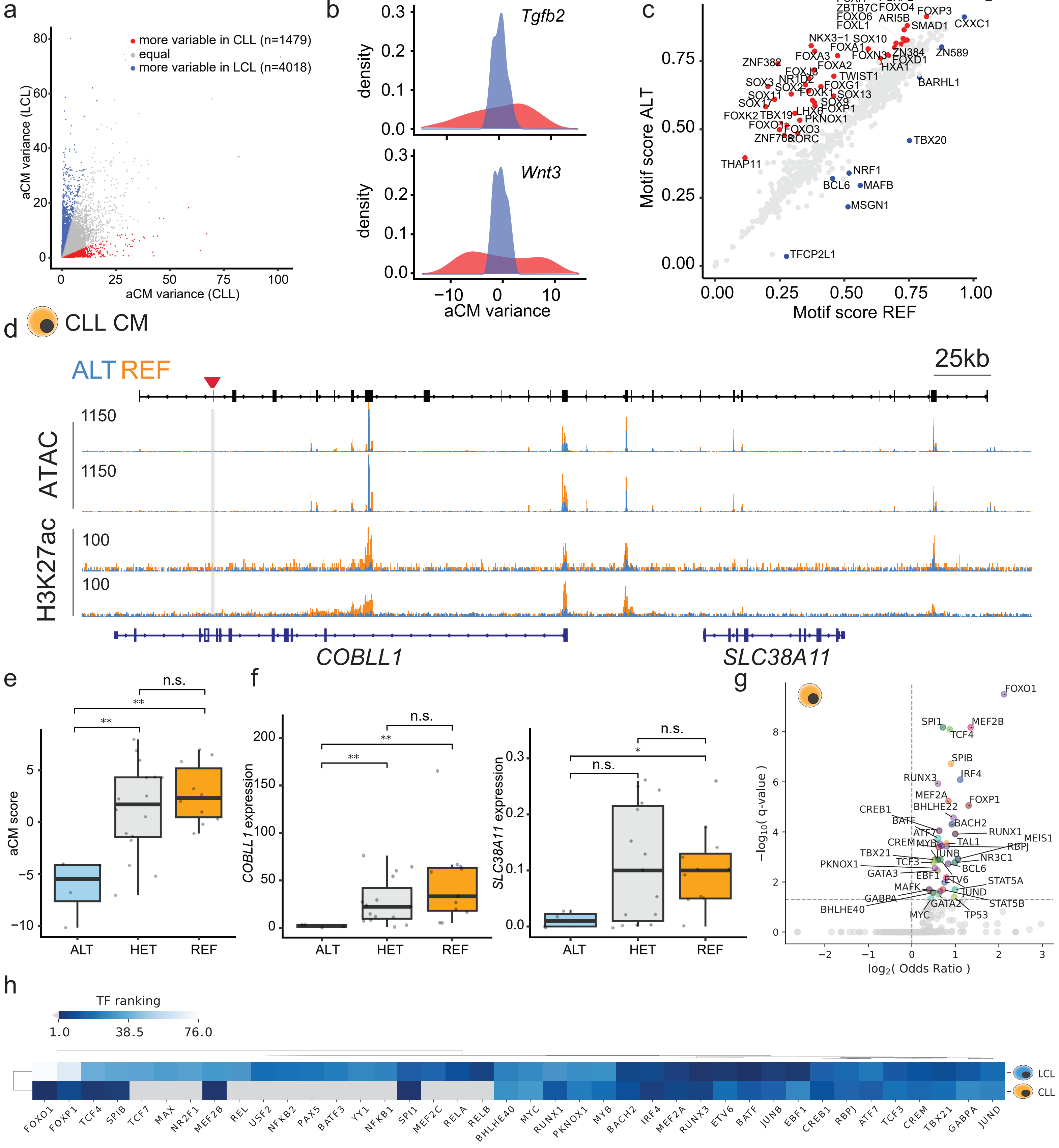

a
more variable in CLL (n=1479)
equal
more variable in LCL (n=4018)
aCM variance (LCL)
aCM variance (CLL)
b
Tgfb2
Wnt3
density
aCM variance
c
Motif score ALT
Motif score REF
FOXI1 FOXP2 ZBTB7C FOXO4 FOXO6 ARI5B FOXP3 FOXL1 SMAD1 CXXC1
NKX3−1 SOX10 FOXA1 ZN384 ZN589 FOXA3 FOXN3 FOXD1 ZNF382 FOXA2 HXA1 FOXJ3 TWIST1 NR1D2 BARHL1 SOX3 SOX2 FOXG1 FOXK1 SOX11 SOX13 SOX9 SOX17 FOXP1 LHX6 FOXK2 TBX19 PKNOX1 FOXO1 FOXO3 ZNF768 RORC TBX20
THAP11 NRF1 BCL6 MAFB MSGN1 TFCP2L1
d
CLL CM
ALT REF
25kb
ATAC
H3K27ac
COBLL1
SLC38A11
e
aCM score
ALT HET REF
n.s.
**
f
COBLL1 expression
SLC38A11 expression
g
−log10( q-value )
log2( Odds Ratio )
FOXO1 SPI1 MEF2B TCF4 SPIB IRF4 RUNX3 MEF2A FOXP1 BHLHE22 CREB1 BATF BACH2 RUNX1 MEIS1 ATF7 CREM MYB TAL1 RBPJ TBX21 JUNB NR3C1 PKNOX1 TCF3 BCL6 GATA3 EBF1 ETV6 STAT5A MAFK JUND GABPA GATA2 STAT5B BHLHE40 MYC TP53
h
TF ranking
1.0 38.5 76.0
LCL
CLL
FOXO1 FOXP1 TCF4 SPIB TCF7 MAX NR2F1 MEF2B REL USF2 NFKB2 PAX5 BATF3 YY1 NFKB1 SPI1 MEF2C RELA RELB BHLHE40 MYC RUNX1 PKNOX1 MYB BACH2 IRF4 MEF2A RUNX3 ETV6 BATF JUNB EBF1 CREB1 RBPJ ATF7 TCF3 CREM TBX21 GABPA JUND

**Figure S5. Distinct CMs are formed in CLL and LCL. a.** Interindividual variance of genomic loci harbouring CMs in either CLL or LCL. Loci that were at least 4 times more variable in either cell type were considered differential. **b.** Example of two regions that show higher interindividual variation in CLL. **c**. Predicted impact of rs895555 on the TF motif score (scaled 0 (no match) to 1 (perfect match)) [102]. **d.** Example depicting the *COBLL1* locus which is induced in a subset of CLL patients. **e.** CM activity stratified by genotype of the highest-ranked candidate associated variant. **f.** Expression of *COBBL1* and *SLC38A11* stratified by genotype of the highest-ranked candidate-associated variant. **g.** Log2 odds ratio versus -log10 q-value of TFBS enrichment within individual cell type when contrasting mapped CM peaks vs simulated CM peaks in CLL (yellow cell). **h.** Heatmap showing the TF ranking for those TFBSs that passed the q-value threshold (0.05) in at least one of the cell types indicated on the right side. Gray color indicates non-significant hits as defined in (**g.**). P-value indications are non-significant (ns) for p-value > 0.05, * for 0.01 < p-value ≤ 0.05, ** for 0.001 < p-value ≤ 0.01.
